# Social steerability modulates perceptual biases

**DOI:** 10.1101/2021.04.28.441710

**Authors:** Hernán Anlló, Katsumi Watanabe, Jérôme Sackur, Vincent de Gardelle

## Abstract

Verbal hints can bias perceptual decision-making, even when the information they provide is false. Whether individuals may be more or less susceptible to such perceptual influences, however, remains unclear. We asked naive participants to indicate the dominant color in a series of stimuli, after giving them a false statement about which color would likely dominate. As anticipated, this statement biased participants’ perception of the dominant color, as shown by a correlated shift of their perceptual decisions, confidence judgments and response times. Crucially, this perceptual bias was more pronounced in participants with higher levels of susceptibility to social influence, as measured by a standard suggestibility scale. Together, these results indicate that even without much apparatus, simple verbal hints can affect our perceptual reality, and that social steerability can determine how much they do so. Susceptibility to suggestion might thus be considered an integral part of perceptual processing.

**Statement of relevance:** At a time when fake news soar, understanding the role that simple verbal descriptions play in how we perceive the world around us is paramount. Extensive research has shown that perception is permeable to well-orchestrated manipulation. Comparatively less attention has been paid to the perceptual impact of false information when the latter is imparted simply and straightforwardly, through short verbal hints and instructions. Here we show that even a single sentence suffices to bias perceptual decision-making, and that critically, this bias varies across individuals as a function of susceptibility to social influence. Considering how here perception was biased by a single, plain sentence, we argue that researchers, communicators and policy-makers should pay careful attention to the role that social suggestibility plays in how we build our perceptual reality.

## Introduction

Will a brief description of an object, true or false, affect the way we see it? Not just our subjective impressions or opinions, but the very contents of our perception. And if so, are we all equally amenable to these description-driven perceptual changes? The challenge of understanding how words can influence our perceptual reality has enthralled philosophers and linguists since times of old, and more recently sparked a widespread interest amongst cognitive scientists, jurists and marketing specialists (Loftus & Palmer, 1974; Cialdini & Goldstein, 2004; Plassman, O’Doherty, Shiv & Rangel, 2008; Falk & Scholz, 2018).

In any instance of communication, descriptive statements can be construed as a form of informational social influence (Deutsch & Gerard, 1955). For example, we might feel inclined to misclassify a line as being longer than another one if we saw other people doing it too (Asch, 1951). During the seminal research on informational social influence, these biases were mainly attributed to an impact of social conformity on decisional processes (Asch, 1951; Asch, 1965; Moscovici & Personnaz, 1980). However, recent evidence has shown that social influence can trigger perceptual judgement distortions through the alteration of sensory and perceptual mechanisms (Mojzisch & Krug, 2008; Germar, Albrecht, Voss & Mojzisch, 2016). Interestingly, these findings are consistent with extensive results of similar nature reported on hypnotic and placebo suggestions, which have also been shown to affect perception in a dramatic fashion, to the point of making participants see things that are not quite there (Spiegel, 2003), preventing them from reading common words (Raz, Kirsch, Pollard & Nitkin-Kaner, 2006), and even blocking them from feeling the pain of a needle going through their skin (Simmons *et al*., 2014). This is in line with theoretical accounts of hypnosis and placebo as settings of social influence, where a researcher or medical professional administers suggestions and prescriptive statements to obtain a perceptual effect, therapeutic or otherwise (Lynn, Kirsch & Hallquist, 2008; Lynn, Laurence & Kirsch, 2015).

One outstanding question is whether all individuals are equally sensitive to this form of social influence. On the one hand, hypnosis research has paid extensive heed to understanding how inter-individual differences in hypnotizability determine the magnitude and limits of suggestion effects. However, the unique social context in which hypnotic and placebo suggestions are delivered, with well-orchestrated manipulations, associative training, or active imagination exercises, render conclusions from their study hard to generalize to more basic contexts of social influence (Terhune et al, 2017; Lynn et al., 2019; Geuter, Koban, Wager, 2017). On the other hand, mainstream informational social influence research has left the matter of inter-individual differences mostly unexplored. Despite early evidence from experimental debriefs (Asch, 1951; Asch, 1965) that only *some* participants ever report experiential changes after being influenced (*i.e.* actually “seeing” the line as longer), identifying the individual differences in how perception can be influenced across neurotypical individuals remains to be developed (Oakley *et al*, 2021).

Traditionally, social psychology inventories of suggestibility are built by assessing our tendency to follow instructions, empathize with others, and comply with a group (Kotov *et al*, 2004). In the present work, we hypothesized that social steerability could predict why some individuals can be influenced into experiencing changes at the perceptual level. To test this hypothesis, we measured the susceptibility to social suggestion of a group of participants with the Short Susceptibility Scale (SSS; Kotov *et al*, 2004), a multidimensional self-administered questionnaire based on introspective judgements and attitudinal behaviors linked to various social influence contexts. These same participants were also asked to indicate the dominant color of a stimulus made of dots of two different colors combined in different proportions, and to report their confidence regarding their perceptual judgment. Crucially, we provided participants with a short written statement about the likelihood of the dominant color for the forthcoming stimuli, at the beginning of each block of trials. This statement was false half of the time, in order to produce a bias that would lead participants to misperceive the dominant color of the stimulus. To minimize effects of compliance or conformity, we prescinded from advisors or confederates, and presented this statement directly along block instructions. Our hypothesis was that our descriptive statement would induce a perceptual change, which would be more pronounced for individuals with higher levels of suggestibility. This led us in turn to enunciate several predictions. First, the false hint would bias perceptual choices relative to a neutral hint, such that participants would overestimate the color mentioned in the hint (hereafter the “target” color). This misrepresentation would be evidenced by a shift (favoring the target color) of the proportion level at which participants deemed both colors as present in equal proportion (*i.e.* the Point of Subjective Equality, hereafter PSE). Second, since we expected subjective task difficulty to be maximal at the PSE, we predicted that minimal confidence ratings and maximal response times would also shift, following the biased PSE. Third, and most crucially, we expected that the inter-individual variability of this bias effect would depend on susceptibility to suggestion, as measured by the SSS. Namely, of all participants, highly suggestible individuals would overestimate the target color the most.

Our results confirmed these shifts in perceptual choices, confidence and response times, as well as their dependence on suggestibility. This suggests that our perceptual reality reflects both sensory inputs and verbal descriptions of these inputs, and that social influence does not require complex suggestion contexts to bias what we see. Considering how social steerability mediated this process despite the absence of acting advisors, normative influence or manipulation techniques, we propose that perceptual decision-making research should consider susceptibility to suggestion as a core feature of perceptual processing.

## Method

We report how we determined our sample size, all data exclusions, all manipulations, and all measures in this study. For further accounts on each of these procedures, see the Supplementary Methods.

### Participants

This experiment was conducted in the Watanabe Cognitive Science Lab (Waseda University, Tokyo, Japan). Acceptable sample sizes (power ≥ 80%) were determined through simulation-based *a-priori* power analyses (see *Supplementary Methods - Section 1*). Participants were recruited throughout the 2018 and 2019 Japanese fiscal year, from the University’s database of volunteers. A total of 56 volunteers were recruited to take part in the experiment. Two participants failed to complete the task, leaving a total of 54 participants (ages between 18 and 23, mean = 19.85 (1.6), 21 female). All volunteers were paid a fixed standard rate of 1000 yen.

### Stimulus and task

On each trial, participants were presented with a stimulus made of two colors, and had to indicate which of the two colors was predominant in the stimulus, as well as their confidence in this perceptual decision. The stimulus was a random-dot kinematogram composed of 100 colored dots (radius = 0.2°, lifetime of 250 ms) presented inside of a circle (radius = 5°) located at the center of the screen. Dots moved without coherence, at a speed of 0.5 deg/s. The stimulus was displayed for a total duration of 1750 ms each trial. In some blocks, the stimulus was made of blue dots and yellow dots. In other blocks, it was made of pink dots and green dots. Nine color proportions were utilized for each color set (between 40% and 60% of the total amount of dots, in steps of 2.5%). Relative luminance, calculated from RGB values as RL = 0.2126R + 0.7152G + 0.0722B, was set at 186 units for all colors. Background color was set at a light shade of grey throughout the entire experiment (R=240, B=240, G=240). Perceptual responses were collected using the arrow keys (the correspondence between color and key was randomly determined for each participant). Confidence ratings were collected by clicking on a scale ranging from 0 (“I am sure I was wrong”) to 50 (“I responded at chance”) to 100 (“I am sure I was correct”). The stimuli and tasks were programmed using Psychtoolbox 3.0.14 (Kleiner *et al.,* 2007) in MATLAB 9.4.0 (R2018a).

### Verbal hints

Trials were presented in blocks, which were preceded by an instruction screen presenting a bias (bias condition) or no bias (control condition). For biased blocks, these instructions stated: “On the following block, there’s a 66% chance that will overcome; namely, it is twice as likely that dots will be more numerous than dots”, where and were replaced by the corresponding color name. Instructions for control blocks stated: “On the following block, there’s a 50% chance that will overcome; namely, it is equally likely for and to outnumber one another”. In order to prevent potential interferences between bias and control blocks, participants received the pink-green set for the control condition and the yellow-blue set for the bias condition, or vice-versa, in a counterbalanced manner across participants. Within each set, the target and distractor colors were also counterbalanced across participants.

### Procedure

Participants were greeted by a research assistant (RA), who asked them to sign a consent form. The RA then proceeded to explain the Type 1 task (the perceptual task) and the Type 2 task (the confidence rating task). The RA insisted on the importance of reflecting upon the confidence ratings even while executing the Type 1 task, and of avoiding extreme or automatic responses that could lead to overestimation or underestimation of confidence levels (*e.g.* to avoid defaulting to 100% or 0% confidence responses if their actual subjective impression was nuanced). Participants were asked to respond as fast as possible to the Type 1 task while remaining precise, and “not to dwell unnecessarily” on the Type 2 task. Participants were informed that by the end of the experiment they would be asked to answer some general questions about themselves.

Each participant was then moved to an individual dim-lit experimental booth, equipped with a low-latency QWERTY keyboard, a mouse, and a chinrest that was fixed 60 centimeters away from a BENQ calibrated color screen. Before starting a practice set, the RA instructed participants to pay careful attention to the instructions that were provided to them at the beginning of each block. Participants then proceeded to do a practice block of 50 trials with a random instruction bias presented at block onset, a random color set and random color ratios. Participants were told that they could stop the training whenever they considered that they had understood the premise of the task (all participants stopped voluntarily before reaching 10 trials). The actual experimental session started immediately afterwards.

Each participant was required to complete 6 randomly-ordered blocks (3 Bias, 3 Control) of trials. Each block presented 90 trials, with 10 trials for each of the 9 proportions of target color in a random order. All blocks began by presenting a specific set of instructions indicating the color set and the false hint about the most likely answer in the case of biased blocks.

Once all blocks were completed, participants were asked to complete Japanese translations of the Short Susceptibility Scale (SSS, excerpted from the Multidimensional Iowa Susceptibility Scale, Kotov *et al*., 2004) and the Ten Item Personality Inventory (TIPI, Benet-Martínez & John, 1998; John & Srivastava, 1999; Gosling, Rentfrow & Swann, 2003). These Japanese translations were translated and then back-translated, and judged as adequate by two Japanese-English bilingual researchers. Scale and item order were randomized for each participant. For each item, participants rated a statement on a scale from false/very unlikely (1) to true/very likely (5 for the SSS, 7 for the TIPI), using the numerical keypad of the keyboard. To prevent suspicions concerning the veracity of the instructional bias, the TIPI and SSS scales were always administered at the end of the task.

Testing sessions lasted between 45 and 65 minutes in total. After the testing session, participants were debriefed on the nature of the experiment. In particular, it was explained to them that task hints for biased blocks were false, and they were asked if they had been suspicious about this fact during the task (none had).

## Analyses

We analyzed our data using generalized linear mixed-models (Agresti, 2002; Jaeger, 2008). To evaluate the significance of a given factor, we compared a model with and a model without this factor using a likelihood ratio test (Pinheiro & Bates, 2000; Bolker, Brooks, Clark, Geange, Poulsen, Stevens & White, 2008; see *Supplementary Methods - Sections 2 & 4*). Fits were performed with the lmer and glmer functions, while convergence estimations and optimizer selection were performed with the allFit function, all from the R package lme4 (Bates, Bolker & Walker, 2015). Once a model was selected, ANOVA tables were computed through Analysis of Deviance (Type II Wald χ² test) to identify significant effects and interactions, and Tukey post-hoc pairwise comparisons were performed when warranted by the results (using car and emmeans R packages; Fox & Weisberg, 2011 and Lenth, 2016, respectively; see *Supplementary Methods - Section 4*).

### Point of subjective equality

Discrimination performance was best represented by the Point of Subjective Equality (PSE). We defined the PSE as the value of target color proportion at which the probability of selecting the target color as the predominant color was equal to 50%. Perceptual decisions were modelled using a probit regression, where the probability of selecting the target color as dominant was a function of the target color proportion, the presence of bias in the block and participants’ susceptibility to suggestion. Given a regression model, we obtained the PSE arithmetically for each condition using the estimated regression coefficients (see *Supplementary Methods - Section 3.1*). To obtain the PSEs’ 95% confidence intervals for each condition, we used a bootstrap approach in which we created 1000 resampled datasets by random sampling with replacement at the trial level (for each condition in each participant), and then refitted the regression model for each of these datasets. Finally, we evaluated the 2.5% and 97.5% quantiles of the distribution of PSEs over these resampled datasets. The same approach was used to evaluate 95% CIs for contrasts between conditions.

### Point of minimal confidence

The point of minimal confidence (PMC) was defined as the value of target color proportion at which confidence levels were at their lowest, for a given condition. Confidence ratings were modelled as a function of target color proportion and its squared value as the explanatory variables, with interactions with Bias and SSS scores. As with the PSE, we used this model’s coefficients to estimate the PMCs arithmetically (see *Supplementary Methods - Section 3.2*), and obtained their 95% CI through bootstrapping, as described above for the PSE.

### Point of maximal response time

The point of maximal response time (PMRT) was defined as the value of target color proportion at which response times were at their highest. As with confidence, response times were modelled as a parabolic function of target color proportion, with additional interactions with Bias and SSS scores (see *Supplementary Materials - Section 3.2*). As before, we used this model’s coefficients to estimate the PMRTs arithmetically, and obtained their 95% CI through bootstrapping.

### Suggestibility

Following previous work (Anlló, Becchio & Sackur, 2017), we considered three categories of suggestibility: High, Medium and Low. To create these categories, we divided total SSS scores by quartiles, and considered the lower quartile to represent Low suggestibility (n=15), the higher quartile High suggestibility (n=16), and the two medium quartiles Medium suggestibility (n=22). Note that these group sizes also reflect the fact that some participants had the same suggestibility scores near the boundaries between quartiles. Our main motivation to create these categories was to be able to obtain group-level estimates of the PSE, PMC and PMRT through fitting, to better illustrate our findings. However, to ensure that none of our effects were artificially generated by this data split, all statistical modelling concerning suggestibility was also performed using alternative models that implemented SSS scores as a continuous variable instead of as a categorical factor. These additional confirmatory analyses supported our findings at every step, and can be found in *Supplementary Materials - Section 4*.

### Additional measures & exclusion criteria

TIPI scores were utilized as a measure of validity for the SSS scale, as means to confirm that SSS scores correlated with personality scores in a similar fashion as in the original SSS norms, and that they described an individual feature distinct from a basic personality trait (Benet-Martínez & John, 1998; John & Srivastava, 1999; Gosling, Rentfrow & Swann, 2003). RTs, on the other hand, were used as a criterion for trial exclusion: we calculated the mean response time for the Type 1 tasks for each level of signal proportion, and excluded trials where RTs were slower or faster than 3 standard deviations of each of the means. A total of 1039 trials across participants were excluded (3.6% of all trials). Additionally, one participant was excluded entirely, for presenting a SSS score higher than 3 standard deviations of the mean. A complete analysis of TIPI scores, as well as comprehensive tests for the metacognitive validity of our confidence measures, can be found in *Supplementary Methods - Sections 4.5 & 4.6*.

## Results

### Perceptual decisions

We anticipated that the probability of reporting one color as dominant in the stimulus would depend on 3 factors: the actual proportion of that color in the stimulus, whether or not that color was mentioned as more likely at the beginning of the block, and participants’ susceptibility to suggestion. To evaluate whether suggestibility was needed to explain perceptual choices, in a generalized mixed model approach, we compared a model that included all 3 factors and their interactions with a simpler version that did not have suggestibility as a factor. The full model outperformed the simpler model (*X*²(4)=14, p<0.01, see *Supplementary Methods - Section 4.1.1*).

To better illustrate the effect of these 3 factors on perceptual choices, we then split participants in High, Medium and Low susceptibility groups (see Figure 2.A *left*), and analyzed the corresponding regression model (see *Supplementary Methods - Section 4.1.2*). As expected, perceptual decisions about color were affected by the color proportion in the stimulus (*X*²(1)=1016, p<0.0001): participants reported seeing the stimulus as “majoritary blue” more often when the proportion of blue dots was larger in the stimulus (the same happened for all colors). Additionally, when a hint towards one color was introduced at the beginning of the block, participants’ decisions were shifted towards that color (*X*²(1)=124, p<0.0001). Crucially, this bias effect also interacted with suggestibility (bias x SSS category interaction: *X*²(2)=37, p<0.0001), confirming our key predictions. These results were replicated by an alternative model, where SSS scores were treated as a continuous predictor instead of a categorical factor, as a measure to confirm the validity of the suggestibility effect (see *Supplementary Methods - Section 4.1.2*).

**Figure 1.**
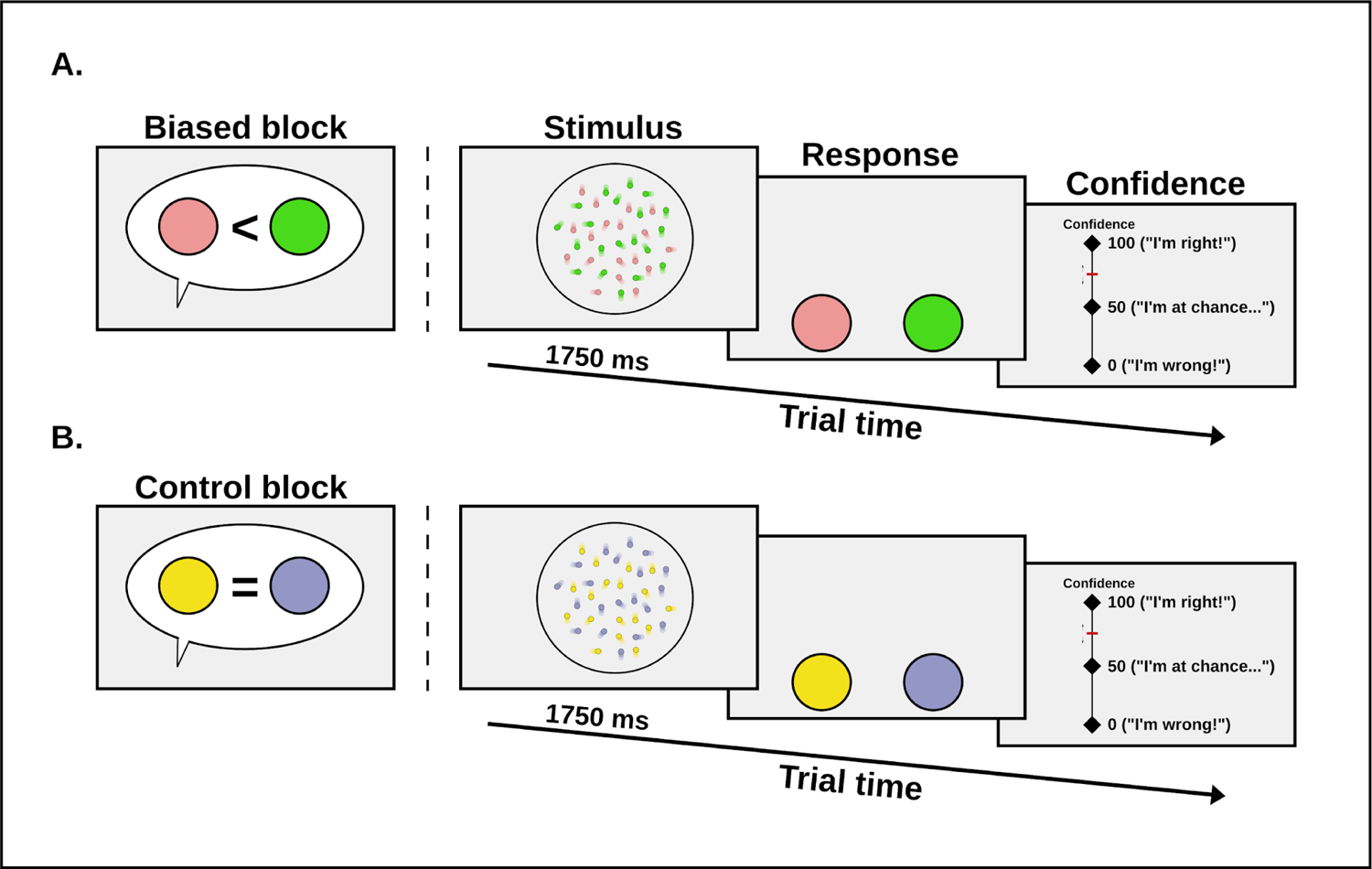
Discrimination paradigm. Participants were presented with a random-dot kinematogram, and then proceeded to give a perceptual discrimination response on which color predominated, followed by a confidence rating (on a scale of 0 to 100). The design was blocked, so that participants did 3 biased blocks and 3 control blocks, for a total of 90 trials per block (total 540 trials). Control and biased blocks used different color sets. **A. Biased blocks.** At the beginning of the block, an instruction defining the bias for the biased blocks was given, explaining there was a 66% chance that dots would overcome dots. **B. Control blocks.** At the beginning of the block, an instruction was given explaining that there was a 50% chance that dots would overcome dots. After stimulus offset, participants had to indicate which color was dominant in that stimulus, and their confidence in this decision.

**Figure 2.**
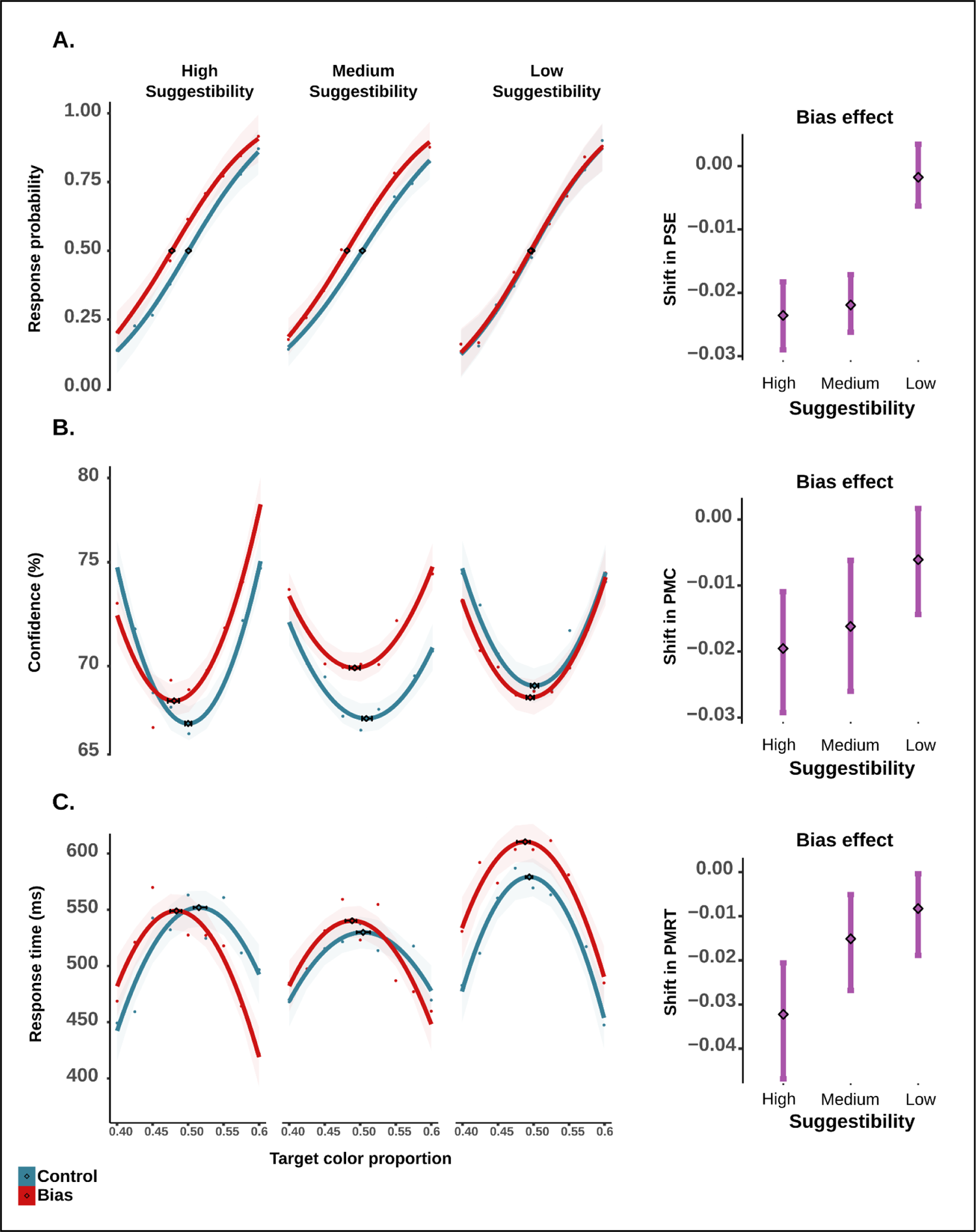
Perceptual choices, confidence and response times. The panels of the left illustrate the variations of perceptual choices (A), of confidence (B), and of response times (C), as a function of the proportion of the target color in the stimulus, separately for high, medium and low suggestibility participants, and for biased and control blocks. Shadowed areas represent the 95% CI of the fit for each condition. The underscored rhomboid dots mark the point of subjective equality (PSE), the point of minimal confidence (PMC), and the point of maximal response times (PMRT) for each condition. Horizontal error bars represent the 95% CI for these estimated values. The panels on the right illustrate the difference between bias and control condition in PSE, PMC, and PMRT, for each level of suggestibility with bars representing 95% CI.

### Point of subjective equality

To examine more closely this interaction, we looked at the point of subjective equality (PSE), that is the value of the stimulus for which participants are indifferent between the two response options. Figure 2.A *(right)* illustrates the difference between biased and control PSEs, together with their 95% CIs, for the 3 subgroups of participants, and shows that participants in the Low suggestibility group exhibited no effect of bias (M=0.002, 95% CI = [−0.006, 0.003]), whereas participants in the Medium and High suggestibility group both did (Medium: M =-0.022 [−0.026, −0.017]; High: M=-0.024 [−0.029, −0.018]). To confirm this difference in the bias effect between the High and Low groups, we computed the 95% CI for that difference and found that it did not include 0 (bias difference 95% CI = [−0.03, −0.01]).

### Confidence ratings

Figure 2.B (*left*) illustrates the confidence with which perceptual decisions are made by participants. As expected, confidence was higher when the color information in the stimulus was more extreme and thus resulted in easier choices. We thus considered a regression model where this could be captured by a quadratic effect of target color proportion on confidence. As was done for perceptual choices, we first confirmed that suggestibility should be included as a factor to account for variations in confidence, as not including it led to a significantly poorer model fit (*X*²(6)=35, p<0.0001, see *Supplementary Methods - Section 4.2.1*).

We then examined these effects in a regression model where confidence was predicted by quadratic and linear effects of target color proportion, together with main effects of bias and suggestibility, and their interactions (see *Supplementary Methods - Section 4.2.2*). The quadratic effect of target color proportion was indeed significant in our data (*X*²(1)=536, p<0.0001), confirming that easier choices were associated with higher confidence. We also conducted further sanity checks showing that confidence also depended on response accuracy and the interaction between difficulty and accuracy (see *Supplementary Methods - Section 4.5*), as typically found in the literature on metaperception (Kepecs, *et al*., 2008; Kepecs & Mainen, 2012).

We also found that confidence was affected by the bias (*X*²(1)=46, p<0.0001), in a way that also interacted with suggestibility (*X*²(2)=58, p<0.0001). This interaction corresponded to the observation that for intermediate target color proportions (*e.g.* at target color proportion=0.5), confidence was higher for biased blocks than for control blocks in participants with High suggestibility (estimate=1.4, SE=0.4, p<0.01) and Medium suggestibility (estimate: 2.5, SE=0.4, p<0.0001) but not for Low suggestibility participants (estimate: −0.4, SE=0.5, p=0.4), as can be seen in the figure. As with perceptual choice, these results were replicated by an alternative model, where SSS scores were treated as a continuous predictor instead of a categorical factor, as a measure to confirm the validity of the suggestibility effect (see *Supplementary Methods - Section 4.2.2*).

### Point of minimal confidence

From the previous fit, we could estimate the point of minimal confidence (PMC), that is, the stimulus value for which decisions are subjectively most uncertain, for all conditions. As shown in figure 2.B *(right)*, this point shifted with the introduction of the biased instruction: confidence was no longer minimal when the stimulus was objectively neutral. Furthermore, this shift in PMCs was present for High suggestibility participants (mean= −0.02, 95% CI = [−0.029, −0.011]), and for Medium suggestibility participants (mean= −0.016 [−0.026, −0.001]) but not for Low suggestibility participants (mean= −0.006 [−0.014, 0.017]). The difference between Low vs High participants was significant, as we found that the 95% CI for this difference in shift did not include 0 [−0.03, −0.002]. This confirmed our prediction that participants incorporated the verbal hint about the stimulus in their confidence ratings too, and even more so when they had high scores on the suggestibility scale.

### Response times

Figure 3.B (*left*) illustrates response times in the perceptual task. As expected, response times changed coherently with task difficulty. As before, we considered a regression model where this could be captured by a quadratic effect of target color proportion on response times. We confirmed that suggestibility improved the prediction of response times variations, as not including it led to a significantly poorer model fit (*X*²(6)=52, p<0.0001, see *Supplementary Methods - Section 4.3.1*). When analyzing the winning model, the effect of task difficulty was confirmed by quadratic and linear effects of target color proportion (*X*²(1)=293, p<0.0001). Further, we observed that response times were also affected by the bias (*X*²(1)=46, p<0.0001), and by the interaction between bias and suggestibility (*X*²(2)=28, p<0.0001). In particular, response times were larger in biased blocks compared to control blocks for Low suggestibility participants (a 32 ms effect for neutral stimuli target color proportion=0.5, SE=10, p<0.01), but not for Medium (10 ms, SE=9, p=0.3) or High suggestibility participants (estimate=-3 ms, SE=10, p=0.8). Complete Analysis of deviance for all regressors (Type 2 Wald *X*² test) and confirmation of these results with a continuous suggestibility score can be found in *Supplementary Methods - Section 4.3.2*.

**Figure 3.**
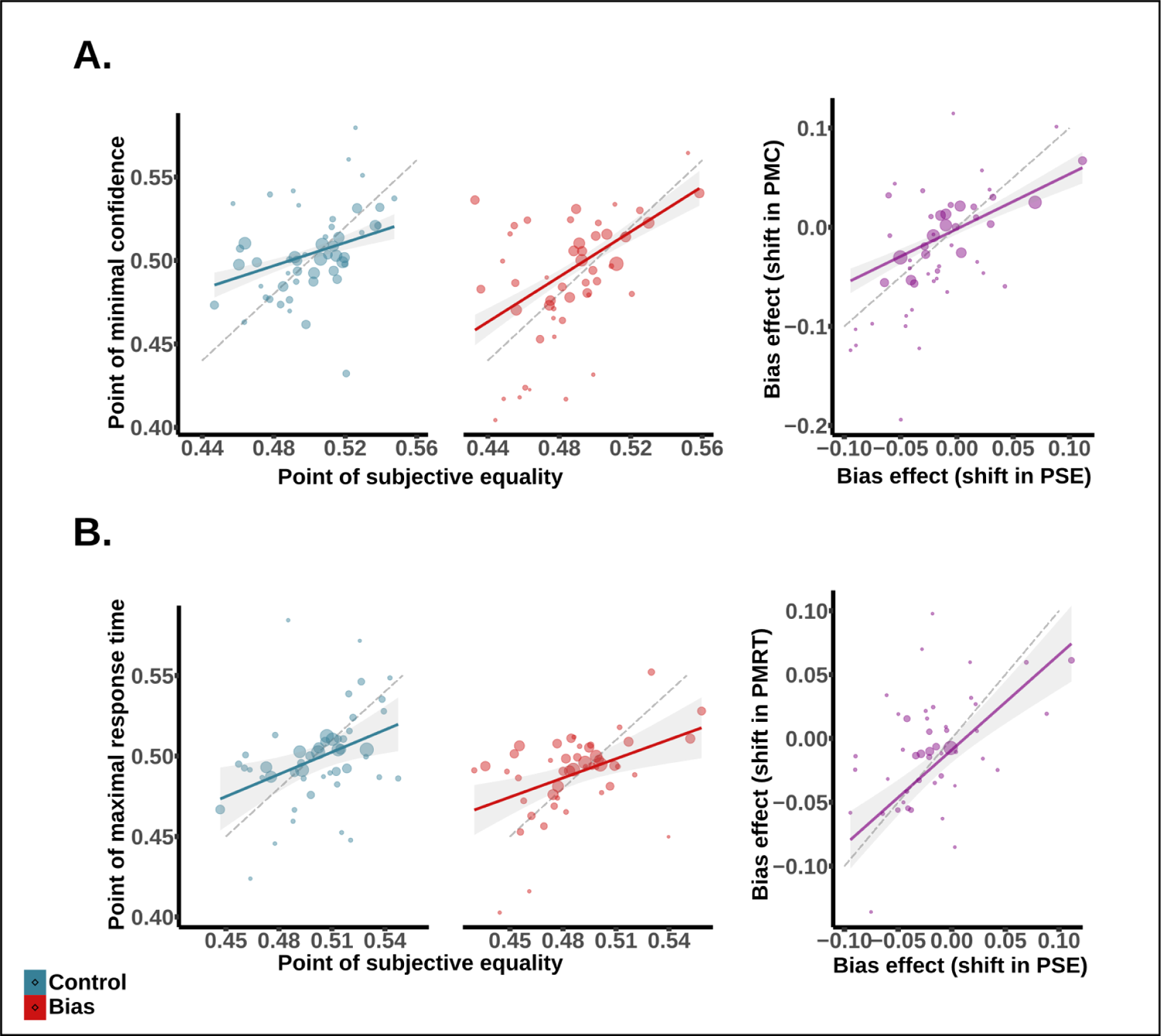
**Joint shifts in performance, confidence and response times. A. Discrimination performance and perceptual confidence. *(Left)*** Regression of individual PSEs against individual PMCs for biased and control conditions. Dot size was determined by the weight of PMCs in the regression, calculated as the inverse of the square sum of biased and control confidence intervals for each condition (*i.e.* the larger the CI, the smaller the weight). This ensured that less noisy individual PMCs contributed more to the regression than noisier PMCs. ***(Right)*** Regression of the shift induced by the bias on PMC (PMC biased minus PMC control) against the same shift for PSE (PSE biased minus PSE control). Dotted line represents the diagonal (slope=1). **B. Discrimination performance and response times *(Left)*** Regression of individual PMRTs against individual PSEs for biased and control conditions. Dot size was determined by the weight of PMRTs in the regression, calculated as the inverse of the PMRT’s 95% CI (*i.e.*, the larger the CI, the smaller the weight). ***(Right)*** Regression of the shift induced by the bias on PMRT (PMRT biased minus PMRT control) against the same shift for PSE (PSE biased minus PSE control).

### Point of maximal response time

From the fitted models, we could estimate the point of maximal response time for each condition. As shown in figure 3.B *(right)*, this point was shifted in biased blocks relative to control blocks. This shift of PMRT was significant in High suggestibility participants (mean= −0.03, 95% CI = [−0.044, −0.02]) and Medium suggestibility participants (mean= −0.016 [−0.029, −0.005]) but not in Low suggestibility participants (mean=-0.006 [−0.015, 0.002]). This difference in shift between Low and High participants was significant (95% CI= [−0.04, −0.009]). These results were in line with what was seen for confidence ratings: subjective choice uncertainty was affected by the verbal hint, particularly in highly suggestible individuals.

### Joint shifts in perceptual decisions, confidence and response times

Results above confirmed that the presence of a simple verbal hint coherently biased perceptual decisions, confidence ratings and response times, and that these shifts were more pronounced for highly susceptible participants. To further examine the relation between these effects, we conducted two additional analyses. First, we tried to predict individual PMCs and PMRTs from individual PSEs. Fitting psychometric curves for confidence and response times was challenging at the individual participant level, and could not be done for some participants (n=3 for the PMC, and n=7 for the PMRT). With the remaining participants, we conducted a weighted regression to predict individual PMCs from PSEs, using the inverse of the PMCs’ 95% CI as individual weights^1^ (figure 3.A, left). Results showed that PSEs significantly predicted PMCs (F_(1,87)_= 21, p<0.0001). The effect of PSEs on PMCs did not interact with bias and suggestibility, when these were included as factors in this regression. In addition, the PSE difference between biased and control also predicted the PMC difference (Figure 3.A, right; F_(1,45)_= 66, p<0.0001). The same relation was confirmed between individual participants’ PSEs and PMRTs, both when predicting PMRT across all conditions (F_(1,84)_= 30, p<0.0001) and when predicting the shift in PMRT between the biased and control condition (F_(1,40)_= 43, p<0.0001). See *Supplementary Methods - Section 4.4* for complete ANOVA tables.

Our second approach was based on comparing whether confidence ratings (or response times) were best predicted at the individual level by the actual stimulus proportion, or by the subjectively biased stimulus proportion. To do so, we fitted two different quadratic models, one based on target color proportion as an explanatory variable, centered around the objective point of equality (*i.e.*, target color proportion minus 0.5), and the other centered around subjective equality (*i.e.*, Target color proportion minus each participant’s PSE). Recentering the proportion this way allowed us to simplify the model (by removing the linear component), and most importantly, to evaluate which was the better predictor of confidence and response times: actual signal equality or subjective equality. Both models also included suggestibility and bias as additional regressors. As expected, the subjective model presented lower AIC and BIC in both cases, indicating that perceptual confidence and response times were better predicted by discrimination performance, rather than actual sensory information.

**Table 1.** Comparison for subjective and objective models predicting confidence and response times

|  | BIC | AIC |
| --- | --- | --- |
| Confidence (Subjective model) | **224351** | **224236** |
| Confidence (Objective model) | 224504 | 224389 |
| Response times (Subjective model) | **224389** | **224306** |
| Response times (Objective model) | 224525 | 224443 |

*Objective models: models where target color proportion was re-centered around objective equality (i.e. 0.5). Subjective models: models where target color proportion was re-centered around each participants' point of subjective equality. BIC: Bayesian Information Criterion. AIC: Akaike Information Criterion. In bold: lower (i. e. better) values.*

## Discussion

In the present work we set out to test whether or not a simple false instruction would bias perceptual decision-making, and to investigate how different individuals would exhibit different sensitivities to this bias. In particular, we hypothesized that the effects of a false preemptive description on perception would depend on suggestibility (construed here as a measure of social steerability). Our findings confirmed these predictions. First, telling participants that a stimulus was more likely to contain more dots of a given color shifted their discrimination responses accordingly, together with their confidence and response times. Second, we confirmed that this perceptual bias varied as a function of suggestibility across participants, and that it was particularly strong for individuals highly susceptible to social influence.

We must discuss first whether the bias induced by our false instruction was perceptual in nature. Indeed, the question of whether perception can be influenced by cognitive factors, including expectations, has been at the heart of intense debates in psychology (see *e.g.* Firestone & Scholl, 2015). Within Signal Detection Theory (Green & Swets, 1966; Macmillan & Creelman, 2005), it has been largely shown that expectations affect perceptual decisions (*e.g.*, Wolfe *et al*., 2005; Ackerman & Landy, 2015; Horowitz, 2017), but whether a shift in responses (as measured by the PSE) reflects a bias in perception or a bias at the decision stage, is hard to know (although see Linares, Aguilar-Lleyda & López-Moliner, 2019 for a suggestion on this matter). In cognitive neuroscience, and particularly within the predictive coding framework, it is commonly assumed that expectations affect perception (see Summerfield & de Lange, 2014; de Lange *et al*., 2018 for reviews), whether these are induced by learned associations between stimuli (Arnal, Morillon, Kell & Giraud, 2009), prior knowledge (Hsieh, 2010) or direct instruction (Summerfield & Koechlin, 2008). Indeed, functional imaging studies have commonly reported effects of prior expectations shaping neural responses to incoming stimuli in perceptual regions, such as the ventral stream (*e.g.* Summerfield & Koechlin, 2008; Pajani *et al*., 2017) or even the primary visual cortex (Kok *et al*., 2013, 2014).

In our study, we relied on confidence ratings and response times to assess the perceptual nature of the bias. Our reasoning was that if participants’ biases reflected a strategic factor (*e.g.* demand effects) instead of true perception, then confidence would be lower and response times higher for biased blocks (compared to control blocks), due to conflicts between the instruction and the information in the stimulus. In addition, in case of a strategic bias, we would anticipate that confidence would remain aligned with the stimulus parameter (*i.e.* lowest for neutral stimuli), and not accompany the PSE shift in suggestible participants. On the contrary, our data showed that confidence was not lower for biased blocks, and that it was centred on the biased point of subjective equality. Similarly, peak response times were also shifted along with the biased PSE. Further, response times were higher for Low suggestibility participants, suggesting that these participants were disturbed by the incongruent instruction, as typically seen in studies portraying cognitive conflict (Cohen, 2014). Together, these observations indicate that the shift in stimulus discrimination likely mirrored the true perceptual experience of participants, rather than being the deployment of a strategy. Interestingly, these findings fall in line with other recent endeavors, where confidence and discrimination criteria were shown to remain coupled despite changes in priors and payoffs (Locke, Gaffin, Hosseinizaveh & Mamassian, 2020).

Our second main result was that the perceptual bias induced by our false description depended on participant suggestibility. The fact that a simple description, even when false, would interact with perception in a manner coherent with susceptibility, reminds us of other forms of influence such as hypnosis and placebo (Geuter, Koban, Wager, 2017; Terhune, Cleeremans, Raz & Lynn, 2017). Indeed, hypnosis and placebo research too have studied manipulations of perception through suggestion, paying special attention to individual differences in susceptibility (Cardeña & Terhune, 2014; Sheiner, Lifshitz & Raz, 2015). Unlike the present study, however, hypnosis and placebo studies rely on the installation of an elaborate social and behavioral context. Active placebo oftentimes couples suggestion with an associative learning phase consisting of convoluted paraphernalia, behavioral routines and elaborate mechanisms of benign deceit (Benedetti *et al*., 2003; Schafer, Colloca & Wager, 2015; Benedetti *et al*. 2016; Geuter, Koban, Wager, 2017). Hypnosis generally uses mental routines to induce participants to actively imagine agency and perceptual changes (Terhune, Cleeremans, Raz & Lynn, 2017). In our work, however, we show that false simple verbal hints and instructions suffice to trigger perceptual modulations, despite the absence of associative learning schemes or mental exercises.

Our findings support the idea of examining all mechanisms of suggestion under the same lens (*i.e.*, as instances of perceptual psychosocial influence), and set the terrain for a study of suggestibility as a core feature of standard perceptual processing. Namely, simple short phrases and verbal instructions may actually condition what we see and feel in everyday life, particularly so if we are highly permeable to social influence. Our findings warrant further study of this phenomenon, and invite the possibility that other findings in the perceptual literature may be the result of suggestion and expectation. For example, recent findings showing the ostensible impact of hypnotizability on the rubber-hand illusion (Lush, Botan, Scott, Seth, Ward & Dienes, 2020), or confederate influence on perceptual judgements during the Asch experiment (Hajnal, Vonk, Zeigler-Hill, 2020) support this proposal.

Along the same lines, our work brings forth the pertinence of inventories where suggestibility is construed as a measure of social steerability to study influence under regular perceptual contexts. Such inventories might be relevant for the development of improved measures of suggestibility, an issue recently revisited in the hypnosis domain (Acunzo & Terhune, 2021; Oakley *et al*, 2021; Kallio, 2021). Moreover, our findings suggest that scales of suggestibility could be valuable tools for studying not only perception, but more generally the construction of judgments, beliefs and trust. Currently, efforts are being made to implement Signal Detection Theory to unravel the appeal of fake news (Batailler, Brannon, Teas & Gawronski, *in press*), and great attention is being paid to the role of direct hints and labels to estimate social media’s trustworthiness and virality (Moravec, Minas & Dennis, 2018). Tackling these questions from a suggestibility perspective appears in our view as a natural step towards the development of a renewed psychology of influence, which should interest researchers, communicators and policy-makers alike.

### Author contributions

HA developed the study concept under the supervision of VdG. JS provided critical feedback. All authors contributed to study design. VdG and HA programmed the experimental task. KW assessed feasibility of the study in Japan. KW and HA performed testing, translation and data collection. VdG, JS and HA prepared a data-analysis plan, which HA executed under the supervision of VdG. JS provided critical feedback on analysis procedures. All authors participated in data interpretation. HA prepared the manuscript under the supervision of VdG, and JS provided critical feedback. All authors approved the final version of the manuscript for submission.

### Author note

This study was supported by grants to HA (17F17008 & 17H00753 from the Japanese Society for the Promotion of Science) and grants to KW (CREST 16817876; Mirai program 20349063, Moonshot Research and Development 20343198 from Japan Science and Technology Agency; KAKENHI 17H06344 & 17H00753, from the Ministry of Education, Culture, Sports, Science and Technology, Japan). The authors declare no conflict of interest. VdG and JS acknowledge the support of Agence Nationale pour la Recherche (Grants ANR-16-CE28-0002, ANR-18-CE28-0015–01, ANR-19-CE28-0019 to VdG and grants ANR-10-LABX-0087 IEC and ANR-10-IDEX-0001-02 PSL to JS).

## Acknowledgements

We thank Koyo Nakamura, Kazuto Toyama and Oishi Hiroyuki for their help with the pilot version, translations of the study scripts and data collection. We would also like to extend our gratitude to all the participants of this study.

## Open practices statements

The data for the experiment is available at the Open Science Framework repository (https://osf.io/cxbaf/?view_only=0b0c4edbd3224aaf845e1a34584505d5). A preprint for this experiment is available online (). Due to a technical error in the preregistration procedure on behalf of the authors, this experiment was not preregistered.

## Supplementary Methods

### 1. Power analyses

In order to estimate statistical power, we chose a simulation-based approach, which is an advantageous technique to use with mixed-models for its flexibility (Green & MacLeod, 2016). The procedure consisted of iterating three consecutive steps: simulating values for the response variable using the model provided, refitting the model to the simulated response, and applying a statistical test to the simulated fit. Since the tested effect was assumed to exist, every positive test was considered a true positive and every negative test was a Type II error. Hence, the power of the test could be calculated from the number of successes and failures at step three.

In the case of our simulated dataset, half of our simulated sample was considered to have a High susceptibility to social Influence, and the other half a Low susceptibility. We simulated one Point of Subjective Equality (PSE) for biased blocks, and one PSE for control blocks. The mean PSE for Lows was the same for biased and unbiased blocks (0.5 ratio of target color, SD=0.01), anticipating that the bias would not affect these individuals (*i.e.*, subjective equality would match on average the objective 0.5 proportion of target color). The mean PSE for Highs was the same as for Lows in the unbiased condition, but shifted to the left for the biased condition by an average of 1 point (*i*. *e*. mean PSE= 0.49 (0.01)). Namely, based on our initial hypothesis, since target color was favored by the bias (*i.e.* “is twice as likely to predominate over”, see *Instructions to Participants*), susceptible participants achieving subjective equality would require less signal than control blocks.

We used the SimR package to conduct these simulations, for sample sizes ranging from 5 to 200 participants (Green & MacLeod, 2016). Alpha was set at 0.05. We considered the model PSE ∼ Bias * SSS category + (1|Participant), and established the detection of a significant Bias x Susceptibility interaction as criterion for success. Each sample was generated, fitted and tested a total of 1000 times, to obtain a power level and the 95% confidence interval for that power level. Figure SM 1 shows power for all samples. Under these conditions, a sample of n=56 yielded a power of 86.2% (95% CI 83.91, 88.28).

**Figure SM.**
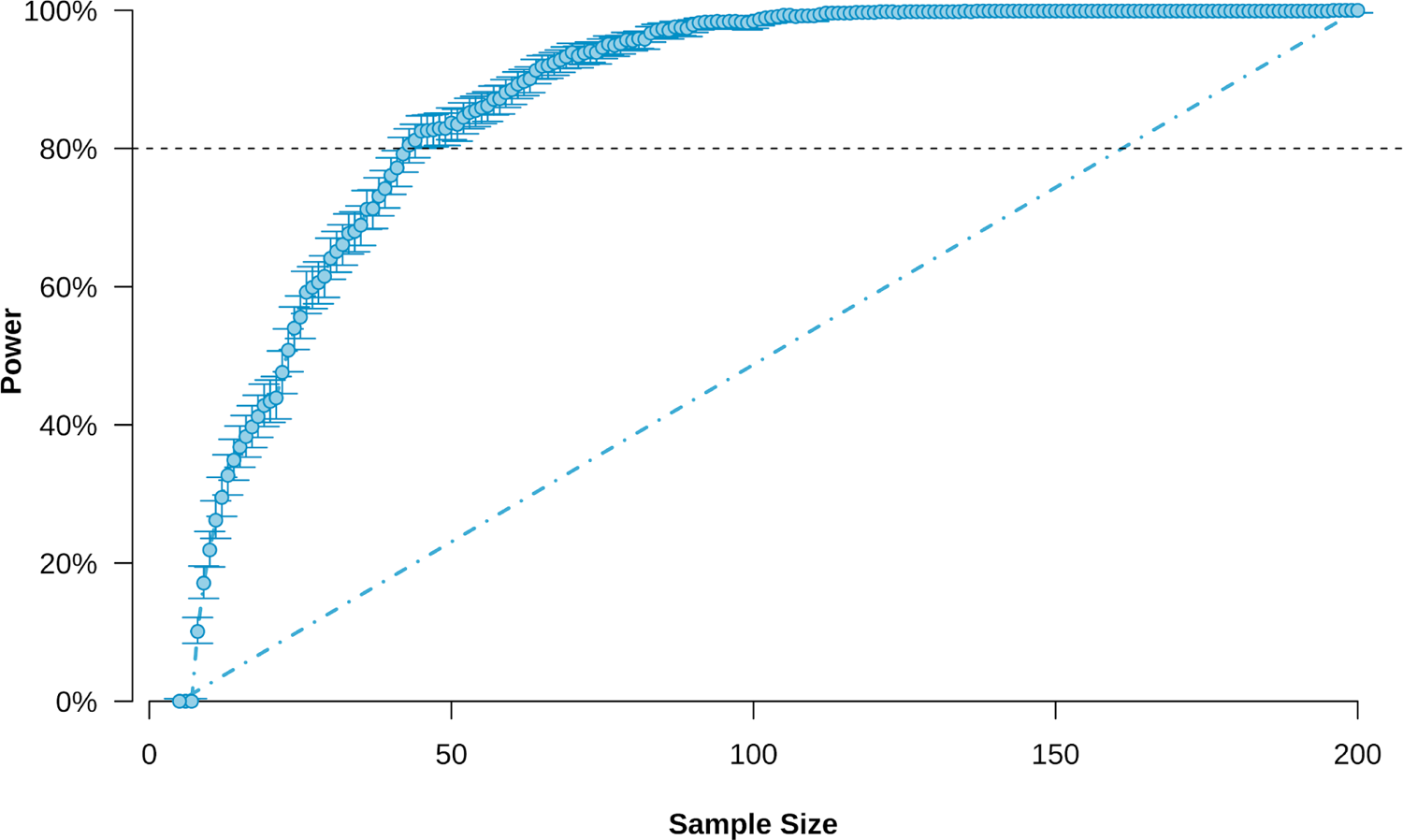
1. Simulation-based power analysis. Power for detecting a significant Bias x suggestibility interaction as a function of sample size (alpha = 0.05). Error bars represent the 95% Confidence Interval for power at each sample. *Black dotted line: 80% power threshold; blue point-dash line: symmetric diagonal*.

### 2. Statistical analyses

#### 2.1 Statistical considerations

We estimated that the best way to analyze the global effects of instructional bias and susceptibility to suggestion without neglecting inter-individual variability and minimizing Type 1 error, was to implement linear mixed-models (Agresti, 2002; Jaeger, 2008). We started by composing a hypothesis-driven full model for each quantity we wanted to predict (response probability, confidence and response time). These models contained target color proportion, type of descriptive hint (biased, control) and suggestibility category (High, Medium, Low) as predictors. Then, we decided whether or not to settle on these hypothesis-driven models by comparing them with simpler models lacking each of these predictors (null models) (Pinheiro & Bates, 2000; Bolker, Brooks, Clark, Geange, Poulsen, Stevens & White, 2008).

In what concerned suggestibility, following previous work (Anlló, Becchio & Sackur, 2017) we considered three categories of suggestibility: High, Medium and Low. To create these categories, we divided total SSS scores by quartiles, and considered the lower quartile to represent Low suggestibility (n=15), the higher quartile High suggestibility (n=16), and the two medium quartiles Medium suggestibility (n=22). Note that these group sizes also reflect the fact that some participants had the same score near the boundaries between quartiles. To ensure that no effects had been artificially generated by this data split, all statistical modelling concerning suggestibility was also performed using alternative models that implemented SSS scores as a continuous variable instead of as a categorical factor. As seen on tables SM4-SM20, these alternative models confirmed our results for every hypothesis-relevant effect and interaction.

For all the models outlined in this work, we coded predictors and variables as follows:

*P(TC)*: probability of selecting target color (response probability)

*TC proportion*: proportion of target color (continuous: between 0.4 and 0.6)

*TC proportion²*: squared proportion of target color.

*Bias*: whether there is a biased hint or a control hint (categorical: Biased, Control)

*SSS category*: suggestibility group (categorical: Low, Medium, High)

*SSS score*: suggestibility raw level obtained from the SSS (continuous: between 21 and 110)

*Confidence*: confidence rating (%)

*RT*: response times (ms)

*Participant*: individual participant tag (used for random intercept allocation)

#### 2.2 Target color identity

The effects of color itself (target color, “TC”: green, pink, yellow, blue) over the probability of selecting the target color (P(TC)) were beyond the interest of this study. Nonetheless, it was important to estimate if the effect of our instructional bias was somewhat attenuated or maximized for a given color. Since the color set (2-level categorical factor: “yellow-blue” / “pink-green”) was covariant with Bias, it was not included as a relevant factor in the main models we evaluated. Instead, in order to evaluate if target color identity had a significant impact on biased responses, we instrumented a target color identity factor between participants to assess the impact of each of the 4 colors in their roles of target and distractor, for both bias and control conditions. We subsetted our data to biased blocks, and compared a model that included this factor with another one that did not. Since the models did not improve by introducing this predictor, it was excluded from the ensuing analysis stages (see *tables SM1-SM3*).

**Table SM1.** Nested model comparisons for target color effects on P(TC) for biased blocks

| | Model | DF | Deviance | $\chi^2$ | P(>$\chi^2$) |
| --- | --- | --- | --- | --- | --- |
| **TC included as factor** | P(TC) ~ TC proportion x SSS score x TC + (TC proportion\|Participant) | 12 | 14823 | 13 | 0.39 |
| **Simplified** | P(TC) ~ TC proportion x SSS score + (TC proportion\|Participant) | -- | 14835 | -- | -- |

**Table SM2.** Nested model comparisons for target color effects on Confidence for biased blocks

| | Model | DF | Deviance | $\chi^2$ | P(>$\chi^2$) |
| --- | --- | --- | --- | --- | --- |
| **TC included as factor** | Confidence ~ (TC proportion + TC proportion<sup>2</sup>) x TC x SSS score + (TC proportion\|Participant) | 18 | 111560 | 25 | 0.12 |
| **Simplified** | Confidence ~ (TC proportion + TC proportion<sup>2</sup>) x SSS score + (TC proportion\|Participant) | -- | 111585 | -- | -- |

**Table SM3.** Nested model comparisons for target color effects on Response Times for biased blocks

| | Model | DF | Deviance | $\chi^2$ | P(>$\chi^2$) |
| --- | --- | --- | --- | --- | --- |
| **TC included as factor** | RT ~ (TC proportion + TC proportion<sup>2</sup>) x TC x SSS score + (TC proportion\|Participant) | 18 | 116162 | 15 | 0.18 |
| **Simplified** | RT ~ (TC proportion + TC proportion<sup>2</sup>) x SSS score + (TC proportion\|Participant) | -- | 116176 | -- | -- |

## 3. PSE, PMC & PMRT computation

We defined the PSE (point of subjective equality) as the value of target color proportion at which the probability of selecting the target color as the predominant color was equal to 50%. As per our hypothesis, we expected the PSE to be smaller for the biased condition, inasmuch as the expectation of “seeing more of the target color” induced through the instruction bias would lead participants to determine equality at lower levels of target color proportion. Further, we expected this difference to be significantly larger as susceptibility to suggestion increased. By the same token, the PMC (point of minimal confidence) was defined as the value of target color proportion at which confidence levels were at their lowest. Namely, we expected confidence to be maximal for near-bound values of target color proportion (*i.e.* trials where the predominant color would be easy to spot), and minimal (around 50%) for trials where the actual proportions of target and distractor were the same (i.e., trials where participants would be inclined to declare themselves closer to chance level). Likewise, we expected the PMC to be smaller for the biased condition (vs. control), and for this difference to significantly increase with susceptibility. Finally, the PMRT (point of maximal response time) was defined as the value of target color proportion at which response times were at their highest. Considering response times as an indicator of processing times, we expected them to increase as the task became harder (*i.e.*, as proportions of target and distractor color dots tended to equality). Here as well, we expected the PMRT to be smaller for the biased condition (vs. control), and for this difference to significantly increase with suggestibility.

Because of the properties of the functions defining their fits, all three of these threshold values could be estimated arithmetically from model coefficients, provided we expressed the linear model formula in the terms of the original fit equations. Further, by bootstrapping the fit and repeating this calculation at each iteration, we were able to estimate the 95% confidence interval for each of these points.

### 3.1 PSE

To assess discrimination performance, we had originally fitted a cumulative Gaussian function (probit) describing the proportion of responses indicating the target color as predominant color. Thanks to its associated link function, the former could be expressed as a linear function:

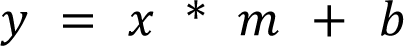

where the distribution function of *y* (in R-base, “pnorm(*y*)”) was the probability of selecting the target color (P(TC)), *x* was the value of the explanatory variable (in our model, TC proportion), *m* was the slope and *b* was the intercept. Considering the link function for the probit fit, P(TC) = 50% when y = 0. Hence

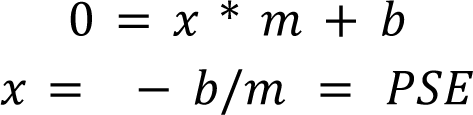

Crucially, *b* and *m* could be obtained from our generalized-mixed model coefficients, for both biased and control conditions in High, Medium and Low susceptibility participants:

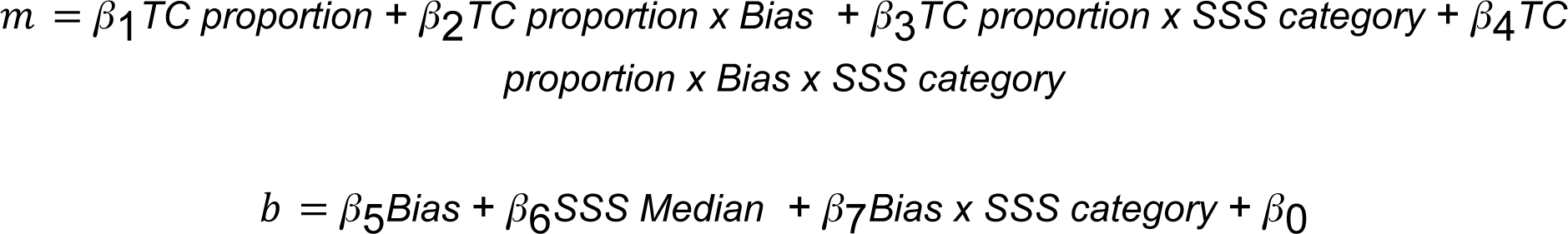

where β_0_ corresponded to the model intercept, and each following β corresponded to the model coefficient for each factor.

### 3.2 PMC & PMRT

The same process was executed to estimate the PMC and the PMRT, given that in the parabolic fit both coincided with the vertex of their parabolas. Given the polynomial function, it follows:

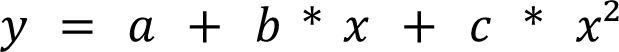

where *y* was the confidence level/response time, *a* was the intercept, and x and *X^2^* were the linear and squared components of the explanatory variable (TC proportion). Since the tangent to the vertex of the parabola had a slope of 0, it was possible to estimate the PMC and the PMRT as

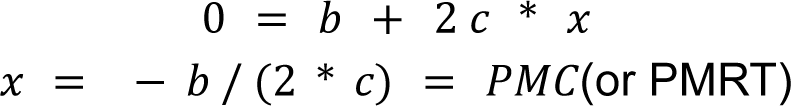

Since *b* and *c* could be obtained from our mixed model coefficients, we executed the calculations as with the PSE above.

## 4. Results

### 4.1 Discrimination performance - perceptual choice

We fitted a cumulative Gaussian model to predict the probability of choosing the target color. Table SM4 shows the hypothesis-driven model, first being compared through maximum-likelihood ratio against a simpler nested alternative that did not include SSS categories. This simplified model was in turn compared to a null model that only included target color proportion. Deviance reduction and chi-square probabilities showed that predictive power increased significantly as hypothesis-relevant factors were included into the model. Hence, we concluded that our original model was better than its alternatives, and was kept for further analysis.

**Table SM4.** Nested model comparisons for discrimination performance, *Suggestibility as a categorical factor (High, Medium, Low)*, *Suggestibility as a continuous predictor (score)*

| | Model | DF | Deviance | $\chi^2$ | P(>$\chi^2$) |
| --- | --- | --- | --- | --- | --- |
| **Hypothesis driven** | P(TC) ~ TC proportion x Bias x SSS category + (TC proportion\|Participant) | 8 | 30556 | 45 | <0.0001 |
| **Simplified** | P(TC) ~ TC proportion x Bias + (TC proportion\|Participant) | 2 | 30601 | 126 | <0.0001 |
| **Null** | P(TC) ~ TC proportion + (TC proportion\|Participant) | -- | 30727 | -- | -- |

*Suggestibility as a continuous predictor (score)*
| | Model | DF | Deviance | $\chi^2$ | P(>$\chi^2$) |
| --- | --- | --- | --- | --- | --- |
| **Hypothesis driven** | P(TC) ~ TC proportion x Bias x SSS score + (TC proportion\|Participant) | 4 | 30583 | 18 | <0.01 |
| **Simplified** | P(TC) ~ TC proportion x Bias + (TC proportion\|Participant) | 2 | 30601 | 126 | <0.0001 |
| **Null** | P(TC) ~ TC proportion + (TC proportion\|Participant) | -- | 30727 | -- | -- |

A summary of the analysis of deviance (Type 2 Wald *X*² test) for the winning model: P(TC) ∼ TC proportion x Bias x SSS score + (TC proportion|Participant), can be found on Table SM5 and Table SM6 below.

**Table SM5.**
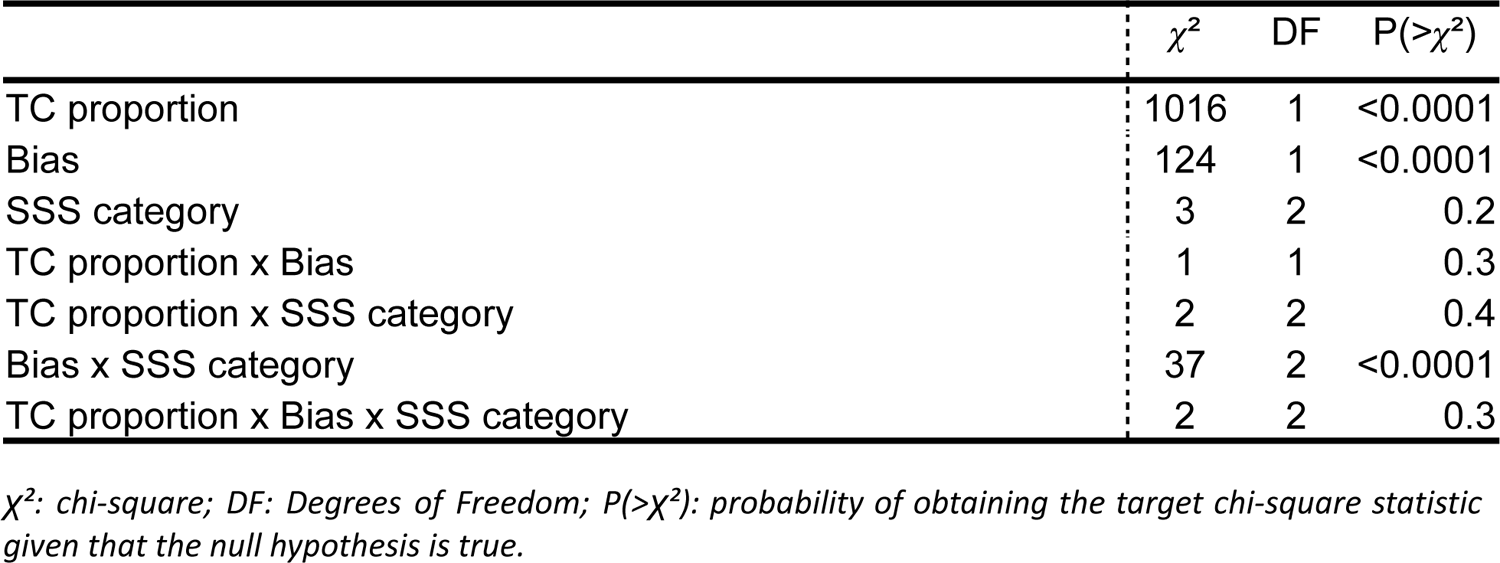
Model P(TC) ∼ TC proportion x Bias x SSS category + (TC proportion|Participant). Analysis of Deviance (Type 2 Wald χ² test) for each effect and interaction.

| | $\chi^2$ | DF | $P(>\chi^2)$ |
| --- | --- | --- | --- |
| TC proportion | 1016 | 1 | <0.0001 |
| Bias | 124 | 1 | <0.0001 |
| SSS category | 3 | 2 | 0.2 |
| TC proportion x Bias | 1 | 1 | 0.3 |
| TC proportion x SSS category | 2 | 2 | 0.4 |
| Bias x SSS category | 37 | 2 | <0.0001 |
| TC proportion x Bias x SSS category | 2 | 2 | 0.3 |

$\chi^2$ : chi-square; DF: Degrees of Freedom; $P(>\chi^2)$ : probability of obtaining the target chi-square statistic given that the null hypothesis is true.

**Table SM6.** Model P(TC) ∼ TC proportion x Bias x SSS score + (TC proportion|Participant). Analysis of Deviance (Type 2 Wald χ² test) for each effect and interaction.

| | $\chi^2$ | DF | $P(>\chi^2)$ |
| --- | --- | --- | --- |
| TC proportion | 974 | 1 | <0.0001 |
| Bias | 125 | 1 | <0.0001 |
| SSS score | 1 | 2 | 0.2 |
| TC proportion x Bias | 1 | 1 | 0.3 |
| TC proportion x SSS score | 1 | 2 | 0.4 |
| Bias x SSS category | 16 | 2 | <0.0001 |
| TC proportion x Bias x SSS category | 0.4 | 2 | 0.5 |

$\chi^2$ : chi-square; DF: Degrees of Freedom; $P(>\chi^2)$ : probability of obtaining the target chi-square statistic given that the null hypothesis is true. Intercept not included on table. SSS scores were rescaled for convergence purposes.

### 4.2 Perceptual confidence

A visual inspection of the proportion of confidence responses selecting target color led us to implement a quadratic fit (confidence was maximal for near-bound values of target color proportion and minimal, around 50%, for trials where the actual proportions of target and distractor were the same). The global fit was then implemented through a mixed-effect model that defined target color proportion (TC proportion) and squared target color proportion (TC proportion**²**) as the explanatory variables, Bias and SSS categories as fixed effects, a random intercept per participant, and random slopes per participant for target color proportion. Table SM7 shows this model being compared through maximum-likelihood ratio with a simpler nested alternative that does not include SSS category. This simplified model was then compared to a null model that only included the linear and squared components of TC proportion. Deviance reduction and chi-square probabilities showed that predictive power increased significantly as hypothesis-relevant factors were included into the model. Hence, we concluded that our original model was better than its alternatives, and it was kept for ensuing analyses.

**Table SM7.** Nested model comparisons for perceptual confidence, Suggestibility as a categorical factor (High, Medium, Low), Suggestibility as a continuous predictor (score)

| | Model | DF | Deviance | $\chi^2$ | P(>$\chi^2$) |
| --- | --- | --- | --- | --- | --- |
| **Hypothesis driven** | Confidence ~ (TC proportion + TC proportion<sup>2</sup>) x Bias x SSS category + (TC proportion\|Participant) | 12 | 224217 | 102 | <0.0001 |
| **Simplified** | Confidence ~ (TC proportion + TC proportion<sup>2</sup>) x Bias + (TC proportion\|Participant) | 3 | 224319 | 79 | <0.0001 |
| **Null** | Confidence ~ TC proportion + TC proportion<sup>2</sup> + (TC proportion\|Participant) | -- | 224398 | -- | -- |

*Suggestibility as a continuous predictor (score)*
| | Model | DF | Deviance | $\chi^2$ | P(>$\chi^2$) |
| --- | --- | --- | --- | --- | --- |
| **Hypothesis driven** | Confidence ~ (TC proportion + TC proportion<sup>2</sup>) x Bias x SSS score + (TC proportion\|Participant) | 6 | 224284 | 35 | <0.0001 |
| **Simplified** | Confidence ~ (TC proportion + TC proportion<sup>2</sup>) x Bias + (TC proportion\|Participant) | 3 | 224319 | 79 | <0.0001 |
| **Null** | Confidence ~ TC proportion + TC proportion<sup>2</sup> + (TC proportion\|Participant) | -- | 224398 | -- | -- |

A summary of the Analysis of Deviance (Type 2 Wald *X*² test) for the winning model: Confidence ∼ (TC proportion + TC proportion²) x Bias x SSS category + (TC proportion|Participant), can be found on Table SM8 and Table SM9 below.

**Table SM8.** Model Confidence ∼ (TC proportion + TC proportion²) x Bias x SSS category + (TC proportion|Participant). Analysis of Deviance (Type 2 Wald χ² test) for each effect and interaction.

| | $\chi^2$ | DF | P(>$\chi^2$) |
| --- | --- | --- | --- |
| TC proportion<sup>2</sup> | 536 | 1 | <0.0001 |
| TC proportion | 519 | 1 | <0.0001 |
| Bias | 46 | 1 | <0.0001 |
| SSS score | 0.3 | 2 | 0.9 |
| TC proportion<sup>2</sup> x Bias | 1 | 1 | 0.3 |
| TC proportion x Bias | 2 | 1 | 0.2 |
| TC proportion<sup>2</sup> x SSS category | 30 | 2 | <0.0001 |
| TC proportion x SSS category | 26 | 2 | <0.0001 |
| Bias x SSS category | 58 | 2 | <0.0001 |
| TC proportion<sup>2</sup> x Bias x SSS category | 1 | 2 | 0.5 |
| TC proportion x Bias x SSS category | 2 | 2 | 0.4 |

$\chi^2$ : chi-square; DF: Degrees of Freedom; P(> $\chi^2$ ): probability of obtaining the target chi-square statistic given that the null hypothesis is true.

**Table SM9.** Model Confidence ∼ (TC proportion + TC proportion²) x Bias x SSS category + (TC proportion|Participant). Analysis of Deviance (Type 2 Wald χ² test) for each effect and interaction.

| | $\chi^2$ | DF | P(>$\chi^2$) |
| --- | --- | --- | --- |
| TC proportion<sup>2</sup> | 534 | 1 | <0.0001 |
| TC proportion | 517 | 1 | <0.0001 |
| Bias | 46 | 1 | <0.0001 |
| SSS score | 0.5 | 1 | 0.5 |
| TC proportion<sup>2</sup> x Bias | 1 | 1 | 0.3 |
| TC proportion x Bias | 2 | 1 | 0.2 |
| TC proportion<sup>2</sup> x SSS score | 14 | 1 | <0.0001 |
| TC proportion x SSS score | 13 | 1 | <0.0001 |
| Bias x SSS score | 10 | 1 | <0.01 |
| TC proportion<sup>2</sup> x Bias x SSS score | 2 | 1 | 0.2 |
| TC proportion x Bias x SSS score | 2 | 1 | 0.1 |

$\chi^2$ : chi-square; DF: Degrees of Freedom; $P(>\chi^2)$ : probability of obtaining the target chi-square statistic given that the null hypothesis is true. Intercept not included on table. SSS scores were rescaled for convergence purposes.

### 4.3 Response times

A visual inspection of the proportion of response time responses when selecting target color led us to implement a quadratic fit (response times were maximal for near-bound values of target color proportion and maximal for trials where the actual proportions of target and distractor color were the same). The global fit was then implemented through a mixed-effect model that defined target color proportion (TC proportion) and squared target color proportion (TC proportion**²**) as the explanatory variables, Bias and SSS categories as fixed effects, a random intercept per participant, and random slopes per participant for target color proportion. Table SM10 shows this model being compared through maximum-likelihood ratio with a simpler nested alternative that does not include SSS category. This simplified model was then compared to a null model that only included the linear and squared components of TC proportion. Deviance reduction and chi-square probabilities showed that predictive power increased significantly as hypothesis-relevant factors were included into the model. Hence, we concluded that our original model was better than its alternatives, and it was kept for ensuing analyses.

**Table SM10.**
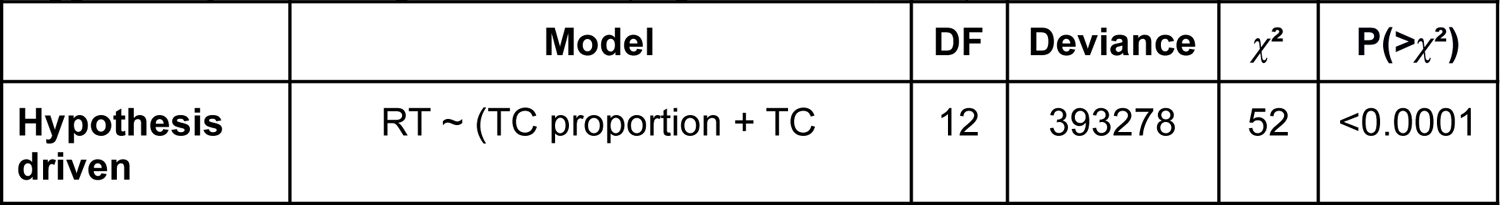
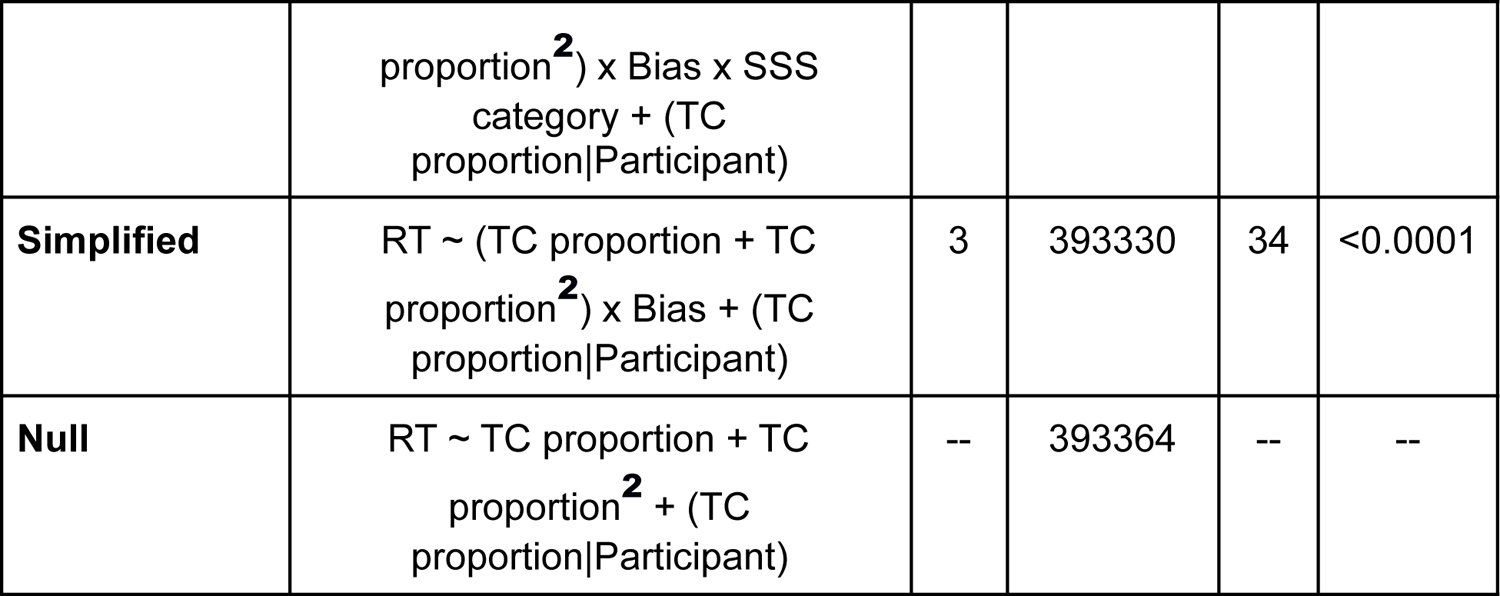
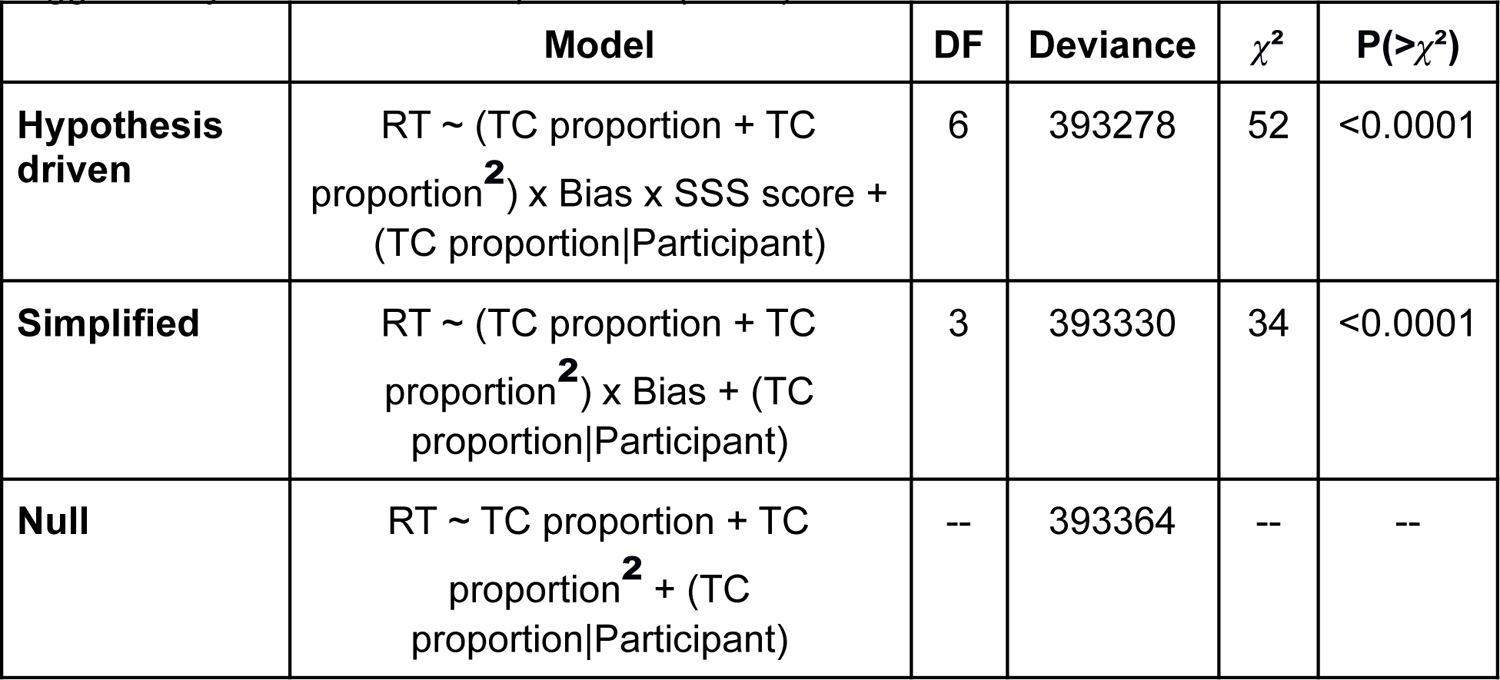
Nested model comparisons for response times, *Suggestibility as a categorical factor (High, Medium, Low)*, *Suggestibility as a continuous predictor (score)*

| | Model | DF | Deviance | $\chi^2$ | $P(>\chi^2)$ |
| --- | --- | --- | --- | --- | --- |
| **Hypothesis driven** | RT ~ (TC proportion + TC proportion<sup>2</sup>) x Bias x SSS category + (TC proportion Participant) | 12 | 393278 | 52 | <0.0001 |
| **Simplified** | RT ~ (TC proportion + TC proportion<sup>2</sup>) x Bias + (TC proportion\|Participant) | 3 | 393330 | 34 | <0.0001 |
| **Null** | RT ~ TC proportion + TC proportion<sup>2</sup> + (TC proportion\|Participant) | -- | 393364 | -- | -- |

*Suggestibility as a continuous predictor (score)*
| | **Model** | **DF** | **Deviance** | $\chi^2$ | **P(>$\chi^2$)** |
| --- | --- | --- | --- | --- | --- |
| **Hypothesis driven** | RT ~ (TC proportion + TC proportion<sup>2</sup>) x Bias x SSS score + (TC proportion\|Participant) | 6 | 393278 | 52 | <0.0001 |
| **Simplified** | RT ~ (TC proportion + TC proportion<sup>2</sup>) x Bias + (TC proportion\|Participant) | 3 | 393330 | 34 | <0.0001 |
| **Null** | RT ~ TC proportion + TC proportion<sup>2</sup> + (TC proportion\|Participant) | -- | 393364 | -- | -- |

A summary of the Analysis of Deviance (Type 2 Wald *X*² test) for the winning model: RT ∼ (TC proportion + TC proportion²) x Bias x SSS category + (TC proportion|Participant), can be found on Table SM12 and SM13 below. While significant interactions of TC proportion x SSS category and TC proportion² x SSS category were found in the model where suggestibility was considered as a categorical factor variable (Table SM11), this was not replicated when suggestibility was taken as a continuous variable (Table SM12). Hence, this interaction was not taken into account.

**Table SM11.** Model RT ∼ (TC proportion + TC proportion²) x Bias x SSS category + (TC proportion|Participant). Analysis of Deviance (Type 2 Wald χ² test) for each effect and interaction.

| | $\chi^2$ | DF | $P(>\chi^2)$ |
| --- | --- | --- | --- |
| TC proportion<sup>2</sup> | 293 | 1 | <0.0001 |
| TC proportion | 285 | 1 | <0.0001 |
| Bias | 7 | 1 | <0.01 |
| SSS score | 0.1 | 2 | 0.9 |
| TC proportion<sup>2</sup> x Bias | 0.4 | 1 | 0.5 |
| TC proportion x Bias | 0.01 | 1 | 0.8 |
| TC proportion<sup>2</sup> x SSS category | 10 | 2 | <0.01 |
| TC proportion x SSS category | 10 | 2 | <0.01 |
| Bias x SSS category | 28 | 2 | <0.0001 |
| TC proportion<sup>2</sup> x Bias x SSS category | 2 | 2 | 0.3 |
| TC proportion x Bias x SSS category | 2 | 2 | 0.4 |

$\chi^2$ : chi-square; DF: Degrees of Freedom; $P(>\chi^2)$ : probability of obtaining the target chi-square statistic given that the null hypothesis is true.

**Table SM12.**
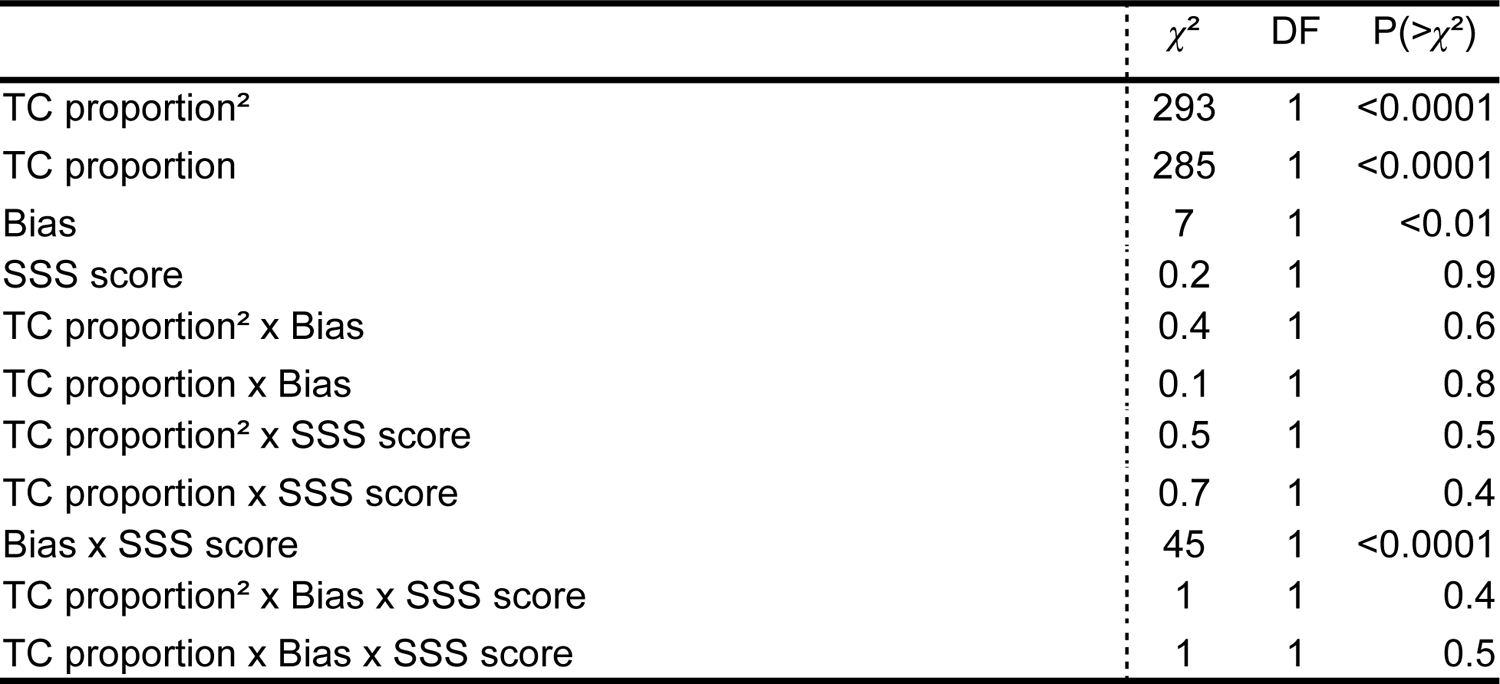
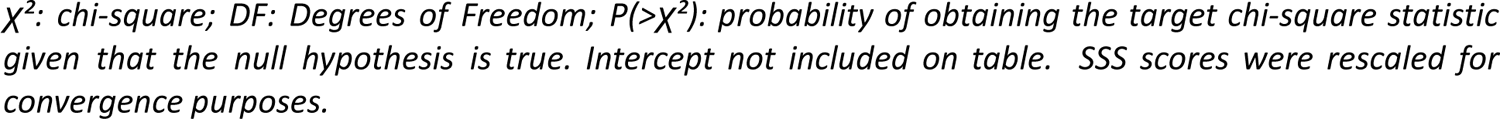
Model RT ∼ (TC proportion + TC proportion²) x Bias x SSS category + (TC proportion|Participant). Analysis of Deviance (Type 2 Wald χ² test) for each effect and interaction.

| | $\chi^2$ | DF | $P(>\chi^2)$ |
| --- | --- | --- | --- |
| TC proportion<sup>2</sup> | 293 | 1 | <0.0001 |
| TC proportion | 285 | 1 | <0.0001 |
| Bias | 7 | 1 | <0.01 |
| SSS score | 0.2 | 1 | 0.9 |
| TC proportion<sup>2</sup> x Bias | 0.4 | 1 | 0.6 |
| TC proportion x Bias | 0.1 | 1 | 0.8 |
| TC proportion<sup>2</sup> x SSS score | 0.5 | 1 | 0.5 |
| TC proportion x SSS score | 0.7 | 1 | 0.4 |
| Bias x SSS score | 45 | 1 | <0.0001 |
| TC proportion<sup>2</sup> x Bias x SSS score | 1 | 1 | 0.4 |
| TC proportion x Bias x SSS score | 1 | 1 | 0.5 |

$\chi^2$ : chi-square; DF: Degrees of Freedom; $P(>\chi^2)$ : probability of obtaining the target chi-square statistic given that the null hypothesis is true. Intercept not included on table. SSS scores were rescaled for convergence purposes.

### 4.4 The performance-confidence & performance-response time links

We fitted the same models as before at the individual participant level, in order to obtain individual PMC, PMRT and PSE values. Individual participants for whom these values could not be estimated through fitting were excluded from this part of the analysis (none for the PSE, n=3 for the PMC, and n=7 for the PMRT). We performed a weighted regression of individual PSEs against individual PMCs and PMRTs for all conditions, using the inverse of the PMCs and PMRTs’ 95% CI respectively, as the weight parameter. Considering the inherent noise of individual-level polynomic fits implemented to obtain PMCs and PMRTs, we took this step to ensure that each PMC and PMRT would affect the regression only as a function of its reliability, in so privileging high quality fits (i.e., the larger the CI, the smaller the weight). We argue that this measure is more reliable and sensitive than taking other measures of goodness of fit, which are limited to establishing a threshold based on observing if the p-value for the difference between fitted model and data is above 0.05.

Tables SM13 and 14 explore the regression of PMCs against PSEs. As shown below, evidence was found only for a significant effect of individual PSEs when regressed against individual PMCs, considering bias and suggestibility. While a significant interaction PSE x SSS category was found for the regression where suggestibility was considered as a categorical factor (Table SM14), this was not replicated when suggestibility was taken as a continuous variable (Table SM15). Hence, this interaction was not taken into account.

**Table SM13.**
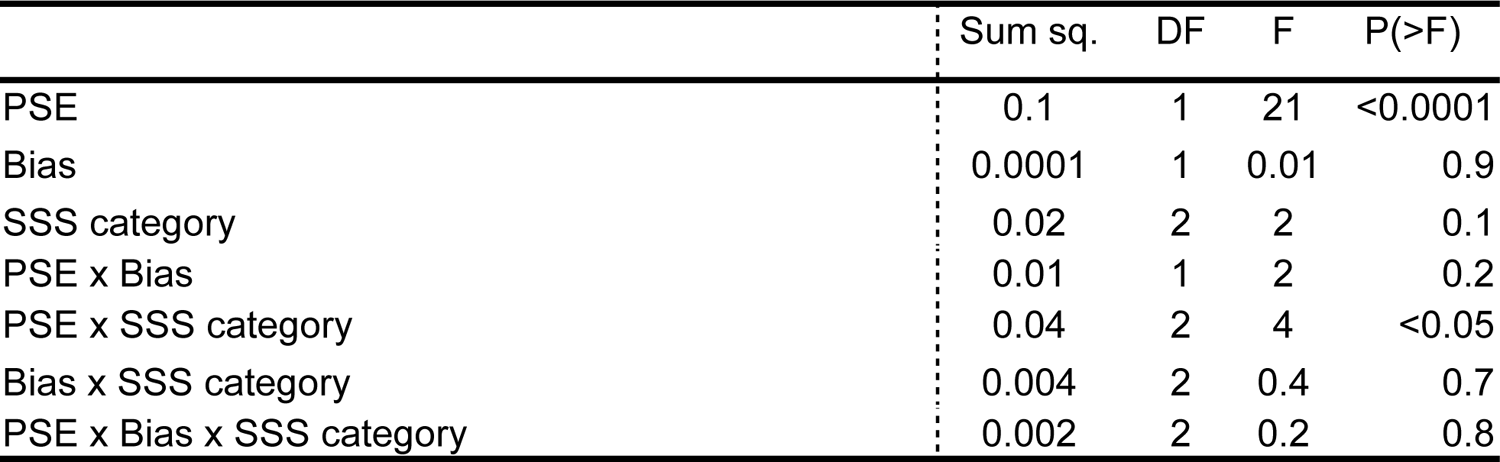
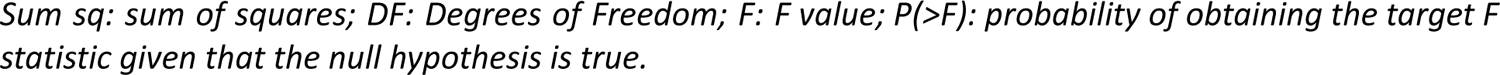
Regression model: PMC ∼ PSE x Bias x SSS category. ANOVA (Type 2) table for each effect and interaction.

|  | Sum sq. | DF | F | P(>F) |
| --- | --- | --- | --- | --- |
| PSE | 0.1 | 1 | 21 | <0.0001 |
| Bias | 0.0001 | 1 | 0.01 | 0.9 |
| SSS category | 0.02 | 2 | 2 | 0.1 |
| PSE x Bias | 0.01 | 1 | 2 | 0.2 |
| PSE x SSS category | 0.04 | 2 | 4 | <0.05 |
| Bias x SSS category | 0.004 | 2 | 0.4 | 0.7 |
| PSE x Bias x SSS category | 0.002 | 2 | 0.2 | 0.8 |

*Sum sq: sum of squares; DF: Degrees of Freedom; F: F value; P(>F): probability of obtaining the target F statistic given that the null hypothesis is true.*

**Table SM14.** Regression model: PMC ∼ PSE x Bias x SSS score. ANOVA (Type 2) table for each effect and interaction.

|  | Sum sq. | DF | F | P(>F) |
| --- | --- | --- | --- | --- |
| PSE | 0.1 | 1 | 19 | <0.0001 |
| Bias | 0.0002 | 1 | 0.04 | 0.8 |
| SSS score | 0.01 | 1 | 2 | 0.2 |
| PSE x Bias | 0.002 | 1 | 0.3 | 0.6 |
| PSE x SSS score | 0.002 | 1 | 0.4 | 0.6 |
| Bias x SSS score | 0.002 | 1 | 0.4 | 0.6 |
| PSE x Bias x SSS score | 0.006 | 1 | 1 | 0.3 |

*Sum sq: sum of squares; DF: Degrees of Freedom; F: F value; P(>F): probability of obtaining the target F statistic given that the null hypothesis is true. SSS scores were rescaled for convergence purposes.*

Anova tables SM15 and SM16 below display the full array of effects and interactions for the models ΔPMC ∼ ΔPSE x SSS category and ΔPMC ∼ ΔPSE x SSS score, respectively.

**Table SM15.**
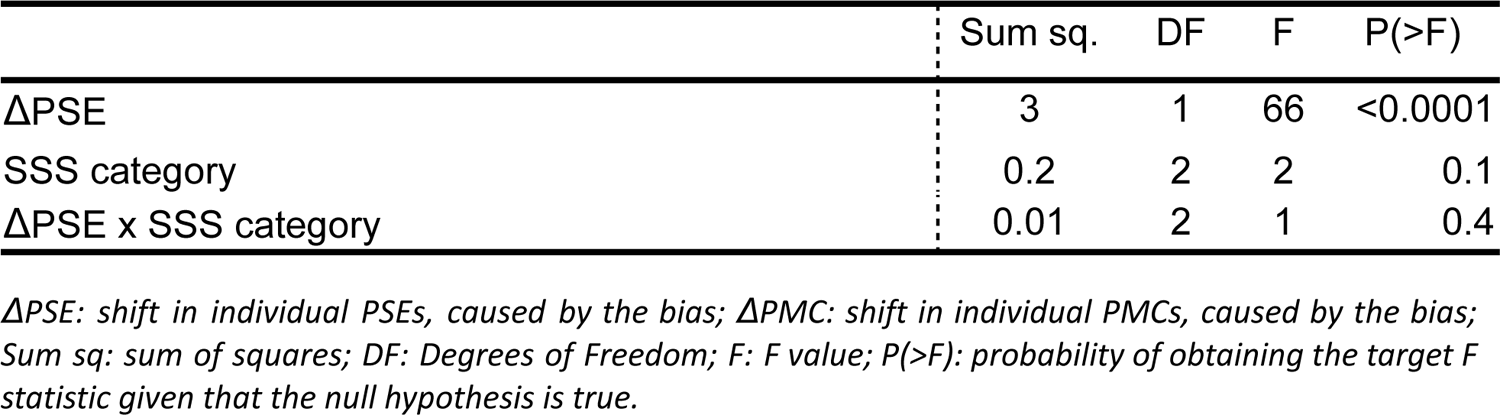
Regression model: ΔPMC ∼ ΔPSE x SSS category. ANOVA (Type 2) table for each effect and interaction.

|  | Sum sq. | DF | F | P(>F) |
| --- | --- | --- | --- | --- |
| $\Delta PSE$ | 3 | 1 | 66 | <0.0001 |
| SSS category | 0.2 | 2 | 2 | 0.1 |
| $\Delta PSE \times SSS \text{ category}$ | 0.01 | 2 | 1 | 0.4 |

*$\Delta PSE$ : shift in individual PSEs, caused by the bias; $\Delta PMC$ : shift in individual PMCs, caused by the bias; Sum sq: sum of squares; DF: Degrees of Freedom; F: F value; P(>F): probability of obtaining the target F statistic given that the null hypothesis is true.*

**Table SM16.**
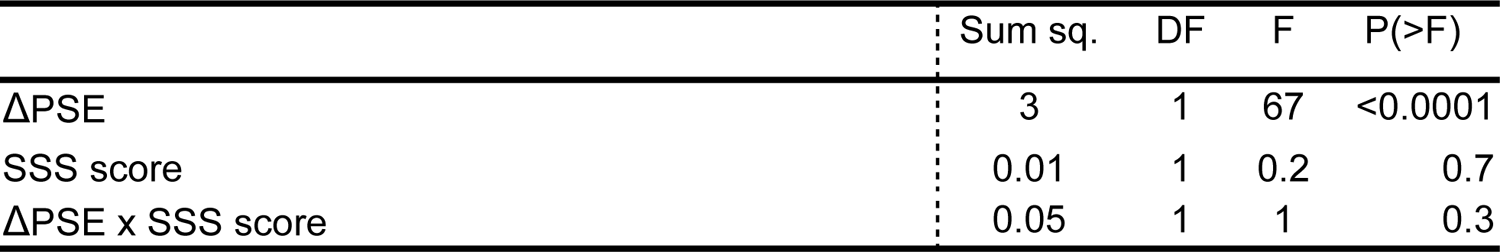
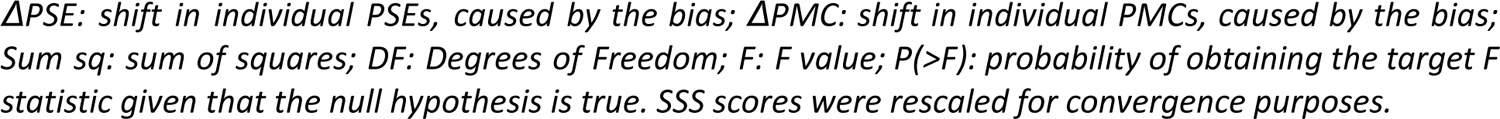
Regression model: ΔPMC ∼ ΔPSE x SSS score. ANOVA (Type 2) table for each effect and interaction.

|  | Sum sq. | DF | F | P(>F) |
| --- | --- | --- | --- | --- |
| $\Delta PSE$ | 3 | 1 | 67 | <0.0001 |
| SSS score | 0.01 | 1 | 0.2 | 0.7 |
| $\Delta PSE \times SSS \text{ score}$ | 0.05 | 1 | 1 | 0.3 |

ΔPSE: shift in individual PSEs, caused by the bias; ΔPMC: shift in individual PMCs, caused by the bias; Sum sq: sum of squares; DF: Degrees of Freedom; F: F value; P(>F): probability of obtaining the target F statistic given that the null hypothesis is true. SSS scores were rescaled for convergence purposes.

Likewise, as shown below on Tables SM17 and SM18 evidence was found only for a significant effect of individual PSEs when regressed against individual PMRTs, considering bias and suggestibility.

**Table SM17.** Regression model: PMRT ∼ PSE x Bias x SSS category. ANOVA (Type 2) table for each effect and interaction.

|  | Sum sq. | DF | F | P(>F) |
| --- | --- | --- | --- | --- |
| PSE | 0.07 | 1 | 30 | <0.0001 |
| Bias | 0.0003 | 1 | 0.1 | 0.7 |
| SSS category | 0.008 | 2 | 2 | 0.2 |
| PSE x Bias | 0.0004 | 1 | 0.2 | 0.7 |
| PSE x SSS category | 0.003 | 2 | 0.7 | 0.5 |
| Bias x SSS category | 0.0005 | 2 | 0.1 | 0.9 |
| PSE x Bias x SSS category | 0.002 | 2 | 0.4 | 0.7 |

Sum sq: sum of squares; DF: Degrees of Freedom; F: F value; P(>F): probability of obtaining the target F statistic given that the null hypothesis is true.

**Table SM18.** Regression model: PMRT ∼ PSE x Bias x SSS score. ANOVA (Type 2) table for each effect and interaction.

|  | Sum sq. | DF | F | P(>F) |
| --- | --- | --- | --- | --- |
| PSE | 0.07 | 1 | 31 | <0.0001 |
| Bias | 0.001 | 1 | 0.6 | 0.5 |
| SSS score | 0.004 | 1 | 1.6 | 0.2 |
| PSE x Bias | 0.0009 | 1 | 0.4 | 0.5 |
| PSE x SSS score | 0.0008 | 1 | 0.3 | 0.6 |
| Bias x SSS score | 0.0005 | 1 | 0.2 | 0.6 |
| PSE x Bias x SSS score | 0.0002 | 1 | 0.1 | 0.8 |

Sum sq: sum of squares; DF: Degrees of Freedom; F: F value; P(>F): probability of obtaining the target F statistic given that the null hypothesis is true. SSS scores were rescaled for convergence purposes.

Anova tables SM19 and SM20 below display the full array of effects and interactions for the models ΔPMRT ∼ ΔPSE x SSS category and ΔPMRT ∼ ΔPSE x SSS score, respectively. While a significant main effect of SSS score was found for the regression where suggestibility was considered as a continuous variable (Table SM20), this was not replicated when suggestibility was taken as a categorical factor (Table SM19). Hence, this interaction was not taken into account.

**Table SM19.** Regression model: ΔPMRT ∼ ΔPSE x SSS category. ANOVA (Type 2) table for each effect and interaction.

|  | Sum sq. | DF | F | P(>F) |
| --- | --- | --- | --- | --- |
| $\Delta PSE$ | 0.7 | 1 | 43 | <0.0001 |
| SSS category | 0.04 | 2 | 1 | 0.3 |
| $\Delta PSE \times SSS \text{ category}$ | 0.01 | 2 | 0.3 | 0.7 |

*$\Delta PSE$ : shift in individual PSEs, caused by the bias; $\Delta PMRT$ : shift in individual PMRTs, caused by the bias; Sum sq: sum of squares; DF: Degrees of Freedom; F: F value; P(>F): probability of obtaining the target F statistic given that the null hypothesis is true.*

**Table SM20.** Regression model: ΔPMRT ∼ ΔPSE x SSS score. ANOVA (Type 2) table for each effect and interaction.

|  | Sum sq. | DF | F | P(>F) |
| --- | --- | --- | --- | --- |
| $\Delta PSE$ | 0.8 | 1 | 56 | <0.0001 |
| SSS score | 0.1 | 1 | 7 | <0.05 |
| $\Delta PSE \times SSS \text{ score}$ | 0.004 | 1 | 0.3 | 0.6 |

*$\Delta PSE$ : shift in individual PSEs, caused by the bias; $\Delta PMRT$ : shift in individual PMRTs, caused by the bias; Sum sq: sum of squares; DF: Degrees of Freedom; F: F value; P(>F): probability of obtaining the target F statistic given that the null hypothesis is true. SSS scores were rescaled for convergence purposes.*

### 4.5 Evaluation of the quality of our confidence measures

Confidence scores constitute a second-order measure. As such, it was important to evaluate if they were informative, and dependent on stimuli parameters, for all experimental conditions. According to Kepecs *et al* (2008; 2012), when plotted as a function of stimulus type and trial outcome, confidence ratings for dominant and non-dominant responses show an opposing V-shaped pattern (or “X pattern”). Conceptually, this means that at each level of stimulus information, participants are more confident of their dominant responses, and less so for their non-dominant responses. As stimulus information decreases and uncertainty increases, confidence ratings for dominant and non-dominant responses approach, only to depart as information increases again. As shown in figure SM3 below, the split of confidence ratings per condition in dominant and non-dominant responses effectively reproduces this pattern. This served as an indication that confidence ratings were coherent and informative, inasmuch as they remained tied to stimulus information, and were generally higher for dominant responses across conditions.

**Figure SM2.**
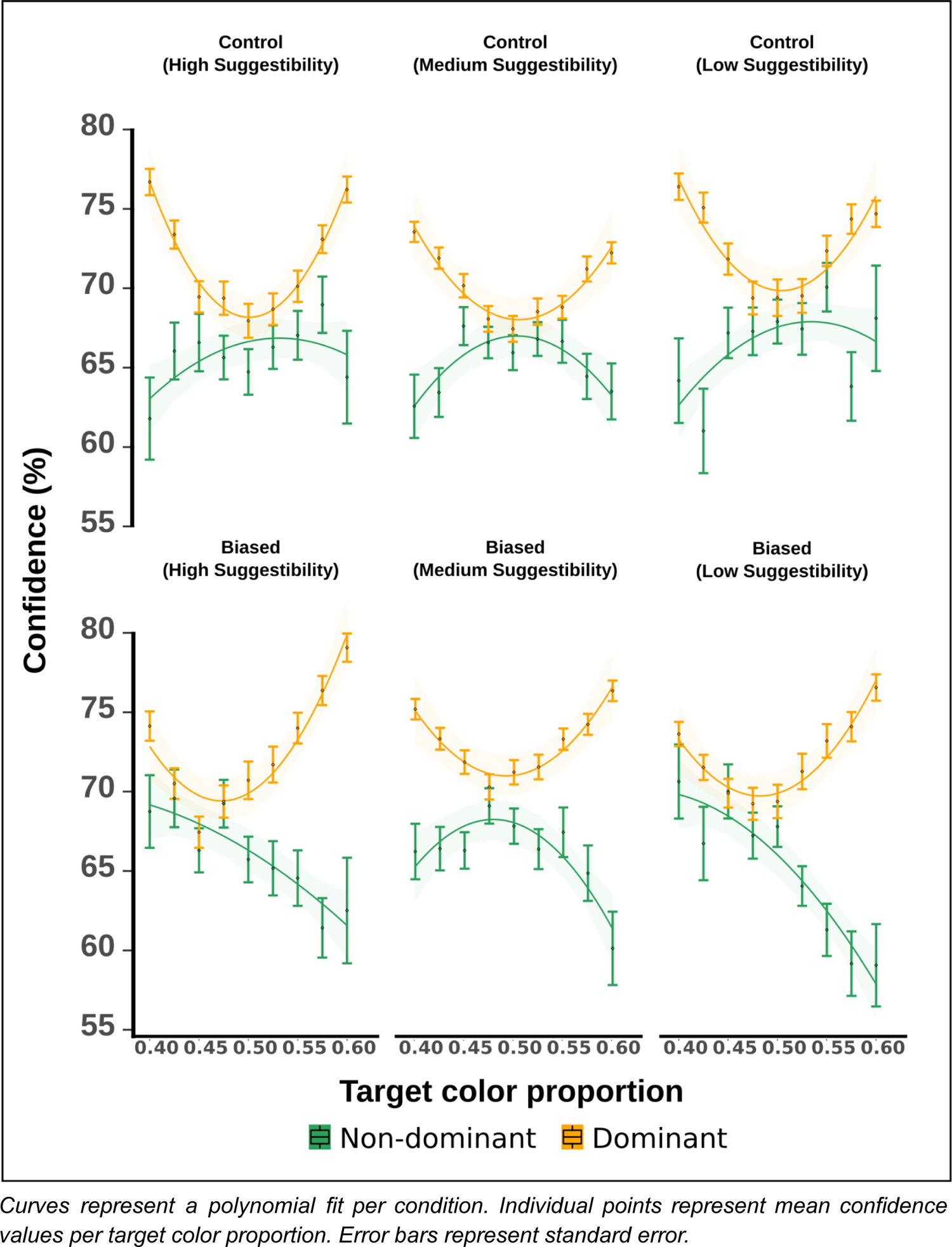
**Confidence ratings for dominant and non-dominant responses.**

### 4.6 TIPI scores

Along with the SSS scores, we also requested participants to complete the Ten Item Personality Inventory (TIPI, Benet-Martínez & John, 1998; John & Srivastava, 1999; Gosling, Rentfrow & Swann, 2003). This inventory consists of a 10-item abridged measurement of the Big Five Personality Inventory. The original SSS score norms (Kotov *et al*, 2004) calculated the Pearson correlation score of Big Five score against suggestibility, in order to determine if the inventory was accurately measuring a distinct individual trait, or a proxy for openness or agreeableness. In the present experiment, asking participants to complete the entire Big Five inventory was not possible due to time constraints, so the TIPI was used instead. While this short inventory is limited in terms of internal consistency and exhibits lower scale-score reliability (Storme *et al*., 2016), it provided us with the possibility of broadly comparing the relationship between suggestibility and personality traits for our sample.

**Table SM21.** On the left the correlations between SSS scores and Big Five personality traits, for the original sample. On the right, the same correlation for our Japanese sample.

|  | SSS (original) | SSS (Japanese sample) |
| --- | --- | --- |
| Extraversion | .11 | -.05 |
| Conscientiousness | -.11 | -.22 |
| Agreeableness | -.04 | -.03 |
| Openness | -.16 | .01 |
| Emotional stability | .22 | -.07 |

Overall, despite some marginal differences between the original and our sample, the low R values show that our translation of the Japanese SSS was also quantifying a different individual psychological trait, not contained by other personality elements.

## Footnotes

1 Considering the inherent noise of individual-level polynomic fits implemented to obtain PMCs and PMRTs, we took this step to ensure that each PMC and PMRT would affect the regression only as a function of its reliability, in so privileging high quality fits (i.e., the larger the CI, the smaller the weight). We argue that this measure is more reliable and sensitive than taking other measures of goodness of fit, which are limited to establishing a threshold based on observing if the p-value for the difference between fitted model and data is above 0.05.

## Notes

### Competing Interest Statement

The authors have declared no competing interest.

## References

1. Ackermann, J. F., & Landy, M. S. (2015). Suboptimal decision criteria are predicted by subjectively weighted probabilities and rewards. Attention, Perception, and Psychophysics, 77(2), 638–658.

2. Acunzo D. & Terhune D. (2021) A Critical Review of Standardized Measures of Hypnotic Suggestibility, International Journal of Clinical and Experimental Hypnosis,69:1, 50–71, DOI: 10.1080/00207144.2021.1833209

3. Agresti, A. (2002) Categorical Data Analysis. Wiley, second edition.

4. Albring A, Wendt L, Benson S, Witzke O, Kribben A, et al. 2012. Placebo effects on the immune response in humans: the role of learning and expectation. PLOS ONE 7(11):e49477

5. Amanzio M, Benedetti F. 1999. Neuropharmacological dissection of placebo analgesia: expectation-activated opioid systems versus conditioning-activated specific subsystems. J. Neurosci. 19(1):484–94

6. Anlló H., Becchio J. & Sackur J. (2017) French Norms for the Harvard Group Scale of Hypnotic Susceptibility, Form A, International Journal of Clinical and Experimental Hypnosis, 65:2, 241–255, DOI: 10.1080/00207144.2017.1276369

7. Arnal L., Morillon B., Kell C., Giraud A (2009) Dual Neural Routing of Visual Facilitation in Speech Processing, Journal of Neuroscience 28 October, 29 (43) 13445-13453; DOI: 10.1523/JNEUROSCI.3194-09.2009

8. Asch SE (1951) Effects of group pressure up on the modification and distortion of judgement. Groups, Leadership and Men, ed Guetzkow H (Carnegie Press, Pittsburgh), pp 177–190.

9. Asch SE (1965) Studies of independence and conformity. Psychol Monogr 70:1–70.

10. Batailler C., Brannon S., Teas P., Gawronski B. (in press) A Signal Detection Approach to Understanding the Identification of Fake News. Perspectives on Psychological Science.

11. Bates, D., Mächler, M., Bolker, B., & Walker, S. (2015). Fitting linear mixed-effects models using lme4. Journal of Statistical Software, 67(1), 1–48. doi:https://doi.org/10.18637/jss.v067.i01

12. Benet-Martínez, V., & John, O. P. (1998). “Los Cinco Grandes” across cultures and ethnic groups: Multitrait-multimethod analyses of the Big Five in Spanish and English. Journal of Personality and Social Psychology, 75, 729–750.

13. Benedetti F, Frisaldi E, Carlino E, Giudetti L, Pampallona A, et al. 2016. Teaching neurons to respond to placebos. J. Physiol. 594(19):5647–60

14. Benedetti F, Pollo A, Lopiano L, Lanotte M, Vighetti S, Rainero I. 2003. Conscious expectation and unconscious conditioning in analgesic, motor, and hormonal placebo/nocebo responses. J. Neurosci. 23(10):4315–23

15. Barton K. (2020), Package MuMIn: Multi-Model Inference [Computer software]. Retrieved from: https://CRAN.R-project.org/package=mumin

16. Bolker B., et al. (2020). GLMM FAQ. Available: https://bbolker.github.io/mixedmodels-misc/glmmFAQ.html

17. Bolker B., Brooks M., Clark C., Geange S., Poulsen j., Stevens H., White J. (2008) Generalized linear mixed models: a practical guide for ecology and evolution Trends in Ecology and Evolution Vol.24 No.3

18. Cialdini R., Goldstein N. (2004) Social Influence: compliance and conformity. Annu. Rev. Psychol. 2004. 55:591–621 doi: 10.1146/annurev.psych.55.090902.142015

19. Cohen M., (2014) A neural microcircuit for cognitive conflict detection and signaling, Trends in Neurosciences, Volume 37, Issue 9, Pages 480-490, ISSN 0166-2236, https://doi.org/10.1016/j.tins.2014.06.004.

20. Colloca L, Petrovic P, Wager TD, Ingvar M, Benedetti F. 2010. How the number of learning trials affects placebo and nocebo responses. Pain 151(2):430–39

21. Fox J., Weisberg S. (2011). An {R} Companion to Applied Regression, Second Edition. Thousand Oaks CA: Sage. URL: http://socserv.socsci.mcmaster.ca/jfox/Books/Companion

22. Cortese A., Amano K., Koizumi A., Kawato M., Lau H. (2016) Multivoxel neurofeedback selectively modulates confidence without changing perceptual performance. Nature Communications | 7:13669 | DOI: 10.1038/ncomms13669

23. de Lange F., Heilbron M., Kok P. (2018) How Do Expectations Shape Perception?, Trends in Cognitive Sciences, Volume 22, Issue 9, Pages 764-779, ISSN 1364-6613, https://doi.org/10.1016/j.tics.2018.06.002.

24. Deutsch, M., & Gerard, H. B. (1955). A study of normative and informational social influences upon individual judgment. The Journal of Abnormal and Social Psychology, 51(3), 629–636. https://doi.org/10.1037/h0046408

25. Dienes, Z., & Hutton, S. (2013). Understanding hypnosis metacognitively: rTMS applied to left DLPFC increases hypnotic suggestibility. Cortex 49, 386–392. doi: 10.1016/j.cortex.2012.07.009

26. Dienes, Z., Perner, J. (2007). Executive control without conscious awareness: The cold control theory of hypnosis. In G. A. Jamieson (Ed.), Hypnosis and conscious states: The cognitive neuroscience perspective (pp. 293–314). Oxford, UK: Oxford University Press.

27. Falk E., Scholz C. (2018) Persuasion, Influence, and Value: Perspectives from Communication and Social Neuroscience. Annu. Rev. Psychol. 2018. 69:18.1–18.28. https://doi.org/10.1146/annurev-psych-122216-011821

28. Firestone, C., Scholl, B.J. Can you *experience* ‘top-down’ effects on perception?: The case of race categories and perceived lightness. Psychon Bull Rev 22, 694–700 (2015). https://doi.org/10.3758/s13423-014-0711-5

29. Germar M, Albrecht T, Voss A, Mojzisch A (2016) Social conformity is due to biased stimulus processing: Electrophysiological and diffusion analyses. Soc Cogn Affect Neurosci 11:1449–1459.

30. Geuter S., Koban L., Wager T. (2017) The Cognitive Neuroscience of Placebo Effects: Concepts, Predictions, and Physiology. Annu. Rev. Neurosci. 2017. 40:167–88. https://doi.org/10.1146/annurev-neuro-072116-031132

31. Gold J, Ding L (2013) How mechanisms of perceptual decision-making affect the psychometric function Progress in Neurobiology 103:98–114. https://doi.org/10.1016/j.pneurobio.2012.05.008

32. Gosling, S. D., Rentfrow, P. J., & Swann, W. B., Jr. (2003). A Very Brief Measure of the Big Five Personality Domains. Journal of Research in Personality, 37, 504–528.

33. Gilman JM, Curran MT, Calderon V, Stoeckel LE, Evins AE (2014) Impulsive Social Influence Increases Impulsive Choices on a Temporal Discounting Task in Young Adults. PLOS ONE 9(7): e101570. https://doi.org/10.1371/journal.pone.0101570

34. Green, P. and MacLeod, C.J. (2016), SIMR: an R package for power analysis of generalized linear mixed models by simulation. Methods Ecol Evol, 7: 493–498. https://doi.org/10.1111/2041-210X.12504

35. Green D, Swets J (1966) Signal Detection Theory and Psychophysics, New York: American Psychological Association.

36. Hajnal A., Vonk J., Zeigler-Hill V. (2020) Peer Influence on Conformity and Confidence in a Perceptual Judgment Task. PSIHOLOGIJA, Vol. 53(1), 101–113 UDC 159.937.24.075 DOI: https://doi.org/10.2298/PSI190107018H

37. Hebart, M. N., Schriever, Y., Donner, T. H. & Haynes, J.-D. (2014) The relationship between perceptual decision variables and confidence in the human brain. Cereb. Cortex 26, 118–130.

38. Horowitz, T. S. (2017). Prevalence in visual search: from the clinic to the lab and back again. Japanese Psychological Research, 59(2),65–108.

39. Hsieh, P.-J. et al. (2010) Recognition alters the spatial pattern of fMRI activation in early retinotopic cortex. J. Neurophysiol. 103, 1501–1507

40. Jaeger, T. (2008). Categorical data analysis: Away from ANOVAs (transformation or not) and towards logit mixed models. Journal of Memory and Language, 59(4):434–446.

41. Jiang, H., White, M. P., Greicius, M. D., Waelde, L. C., & Spiegel, D. (2016). Brain activity and functional connectivity associated with hypnosis. Cerebral Cortex. doi: 10.1093/cercor/bhw220

42. John, O. P & Srivastava, S. (1999). The Big Five trait taxonomy: History, measurement, and theoretical perspectives. In L. A. Pervin, & O. P. John (Eds.), Handbook of personality: Theory and research(pp.102–138). New York: Guilford Press.

43. Kallio S. (2021) Time to update our suggestibility scales. Consciousness and Cognition 90, 103103. https://doi.org/10.1016/j.concog.2021.103103

44. Kepecs A, Mainen ZF. (2012) A computational framework for the study of confidence in humans and animals. Philosophical Transactions of the Royal Society of London. Series B, Biological Sciences. 367: 1322–37. PMID 22492750 DOI: 10.1098/rstb.2012.0037

45. Kepecs, A., Uchida, N., Zariwala, H. A. & Mainen, Z. F. (2008) Neural correlates, computation and behavioural impact of decision confidence. Nature 455,227–231.

46. Kiani, R. & Shadlen, M. N. (2009) Representation of confidence associated with a decision by neurons in the parietal cortex. Science 324, 759–764.

47. Kok P., Brouwer G., van Gerven M., de Lange F. (2013) Prior Expectations Bias Sensory Representations in Visual Cortex. Journal of Neuroscience, October 9 33(41):16275–16284 • 16275

48. Konishi, M., Compain, C., Berberian, B. et al. (2020) Resilience of perceptual metacognition in a dual-task paradigm. Psychon Bull Rev. https://doi.org/10.3758/s13423-020-01779-8

49. Kotov RI, Bellman SB, Watson DB. Multidimensional Iowa Suggestibility Scale (MISS) Brief Manual[Internet]. 2004 [cited 1 Apr 2015] pp. 7–10. Available: http://medicine.stonybrookmedicine.edu/system/files/MISSBriefManual.pdf

50. Landry M., Appourchaux K., Raz A. (2014) Elucidating unconscious processing with instrumental hypnosis. Frontiers in Psychology. 28 July, doi: 10.3389/fpsyg.2014.00785

51. Lenth R. (2020). Estimated Marginal Means, aka Least-Squares Means [Computer software]. Retrieved from https://CRAN.R-project.org/package=emmeans

52. Lifshitz M, Bonn Aubert N, Kashem IF, Raz A (2013) Using suggestion to modulate automatic processes: From Stroop to McGurk and beyond. Cortex 49: 463–473. doi:10.1016/j.cortex.2012.08.007. PubMed: 23040173.

53. Linares, D., Aguilar-Lleyda D. & López-Moliner J. (2019) Decoupling sensory from decisional choice biases in perceptual decision-making. eLife 2019;8:e43994, https://doi.org/10.7554/eLife.43994.001

54. Linares, D., & López-Moliner, J. (2016). quickpsy: An R package to fit psychometric functions for multiple groups. The R Journal, 8(1), 122–131.

55. Locke S., Gaffin Cahn E., Hosseinizaveh N, Mamassian P., Landy M (2020) Priors and payoffs in confidence judgments. Attention, Perception, & Psychophysics. https://doi.org/10.3758/s13414-020-02018-x

56. Loftus E., Palmer J. (1974) Reconstruction of Automobile Destruction: An Example of the Interaction Between Language and Memory. Journal of verbal learning and verbal behavior, 13, 585–589. https://doi.org/10.1016/S0022-5371(74)80011-3

57. Lush P., Botan V., Scott R., Seth A., Ward J., Dienes Z. (2020) Trait phenomenological control predicts experience of mirror synaesthesia and the rubber hand illusion. Nature communications |11:4853 | https://doi.org/10.1038/s41467-020-18591

58. Lynn, S. J., Green, J. P., Polizzi, C. P., Ellenberg, S., Gautam, A., & Aksen, D. (2019). Hypnosis, hypnotic phenomena, and hypnotic responsiveness: Clinical and Research Foundations-A 40-Year Perspective. International Journal of Clinical and Experimental Hypnosis, 67(4), 475–511. doi:10.1080/00207144.2019.1649541

59. Lynn, S.J., Kirsch, I., Hallquist, M.N., (2008) Social cognitive theories of hypnosis. In: Nash, M.R., Barnier, A.J. (Eds.), The Oxford Handbook of Hypnosis: Theory, Research and Practice. Oxford University Press Oxford, Oxford, pp. 111–139

60. Lynn, S. J., Laurence, J. R., & Kirsch, I. (2015). Hypnosis, suggestion, and suggestibility: an integrative model. American Journal of Clinical Hypnosis, 57(3), 314–329. doi:10.1080/00029157.2014.976783

61. Magalhães De Saldanha da Gama PA, Slama H, Caspar EA, Gevers W, Cleeremans A (2013) Placebo-Suggestion Modulates Conflict Resolution in the Stroop Task. PLoS ONE 8(10): e75701. doi:10.1371/journal.pone.0075701

62. Macmillan, N. A., & Creelman, C. D. (2005). *Detection theory: A user’s guide* (2nd ed.). Lawrence Erlbaum Associates Publishers.

63. McGeown, W. J., Venneri, A., Kirsch, I., Nocetti, L., Roberts, K., Foan, L., et al (2012). Suggested visual hallucination without hypnosis enhances activity in visual areas of the brain. Consciousness and Cognition, 21(1), 100–116.

64. Mojzisch A, Krug K (2008) Cells, circuits, and choices: Social influences on perceptual decision making. Cogn Affect Behav Neurosci 8:498–508.

65. Montgomery GH, Kirsch I. 1997. Classical conditioning and the placebo effect. Pain 72(1–2):107–13

66. Moravec P, Minas R., Dennis A. (2018) Fake News on Social Media: People Believe What They Want to Believe When it Makes No Sense at All. Kelley School of Business Research Paper No. 18-87, Available at SSRN: https://ssrn.com/abstract=3269541 or http://dx.doi.org/10.2139/ssrn.3269541

67. Moscovici S, Personnaz B (1980) Minority influence and conversion behaviour in a perceptual task. J Exp Soc Psychol 16:270–282.

68. Pajani, A., Kouider, S., Roux, P., de Gardelle V. (2017) Unsuppressible Repetition Suppression and exemplar-specific Expectation Suppression in the Fusiform Face Area. Sci Rep 7, 160 (2017). https://doi.org/10.1038/s41598-017-00243-3

69. Pinheiro J.C., Bates D.M. (2000). Mixed-Effects Models in S and SPLUS. New York: Springer.

70. Plassmann H., O’Doherty J., Shiv B., Rangel A. (2008) Marketing actions can modulate neural representations of experienced pleasantness. PNAS January 22, 105 (3) 1050-1054; https://doi.org/10.1073/pnas.0706929105

71. Price DD, Milling LS, Kirsch I, Duff A, Montgomery GH, Nicholls SS. (1999) An analysis of factors that contribute to the magnitude of placebo analgesia in an experimental paradigm. Pain 83:147–56

72. Oakley, A., Walsh, E., Mehta, M. A., Halligan, P. W., & Deeley, Q. (2021). Direct verbal suggestibility: Measurement and significance. Consciousness and Cognition, 89, 1–20. https://doi.org/10.1016/j.concog.2020.103036

73. R Core Team. (2018). R: A language and environment for statistical computing [Computer software]. Vienna, Austria: R Foundation for Statistical Computing. Retrieved from https://www.R-project.org/

74. Rahnev D, Lau H, de Lange FP (2011) Prior expectation modulates the interaction between sensory and prefrontal regions in the human brain. J Neurosci. 2011 Jul 20;31(29):10741-8. doi: 10.1523/JNEUROSCI.1478-11.2011. PMID: 21775617; PMCID: PMC6622631.

75. Raz, A., Kirsch, I., Pollard, J., & Nitkin-Kaner, Y. (2006). Suggestion reduces the Stroop effect. Psychological Science, 17, 91–95. https://doi.org/10.1111/j.1467-9280.2006.01669.x

76. Schafer SM, Colloca L, Wager TD. 2015. Conditioned placebo analgesia persists when subjects know they are receiving a placebo. J. Pain 16(5):412–20

77. Sheiner, E. O., Lifshitz, M., & Raz, A. (2015, November 30). Placebo Response Correlates With Hypnotic Suggestibility. Psychology of Consciousness: Theory, Research, and Practice. Advance online publication. http://dx.doi.org/10.1037/cns0000074

78. Simmons K., Ortiz R., Kossowsky J., Krummenacher P., Grillon C., Pine D., Colloca L.,Pain and placebo in pediatrics: A comprehensive review of laboratory and clinical findings, PAIN (2014), doi: http://dx.doi.org/10.1016/j.pain.2014.08.036

79. Spiegel D. (2003) Negative and Positive Visual Hypnotic Hallucinations:Attending Inside and Out, International Journal of Clinical and Experimental Hypnosis, 51:2,130–146, DOI: 10.1076/iceh.51.2.130.14612

80. Summerfield C., Koechlin E., (2008) A Neural Representation of Prior Information during Perceptual Inference, Neuron, Volume 59, Issue 2, Pages 336-347, ISSN 0896-6273, https://doi.org/10.1016/j.neuron.2008.05.021.

81. Summerfield, C., de Lange, F. (2014) Expectation in perceptual decision making: neural and computational mechanisms. Nat Rev Neurosci 15, 745–756. https://doi.org/10.1038/nrn3838

82. Terhune D. B., Cleeremans A., Raz A., Lynn S.J. (2017) Hypnosis and top-down regulation of consciousness. Neuroscience & Behavioral Reviews. doi: http://dx.doi.org/10.1016/j.neubiorev.2017.02.002

83. Terhune, D. B., Polito, V., Barnier, A. J., & Woody, E. Z. (2016) Variations in the Sense of Agency During Hypnotic Responding: Insights From Latent Profile Analysis. Psychology of Consciousness: Theory, Research, and Practice. Advance online publication. http://dx.doi.org/10.1037/cns0000107

84. Teufel C., Dakin S., Fletcher P. (2018) Prior object-knowledge sharpens properties of early visual feature-detectors. Scientific Reports | 8:10853 | DOI:10.1038/s41598-018-28845-5

85. Vandenbroucke A., Fahrenfort J., Meuwese J., Scholte H., Lamme V. (2014) Prior Knowledge about Objects Determines Neural Color Representation in Human Visual Cortex. Cerebral Cortex; 26:1401–1408 doi:10.1093/cercor/bhu224

86. Wager T., Atlas L. (2015) The neuroscience of placebo effects: connecting context, learning and health. Nature Reviews | Neuroscience. VOLUME 16. doi:10.1038/nrn3976

87. Wickham, H. (2017). Tidyverse: Easily install and load the “tidyverse” [Computer software]. Retrieved from https://CRAN.R-project.org/package=tidyverse

88. Wolfe, J. M., Horowitz, T. S., & Kenner, N. M. (2005). Rare items often missed in visual searches. Nature, 435, 439–440.

## References

89. Benet-Martínez, V., & John, O. P. (1998). “Los Cinco Grandes” across cultures and ethnic groups: Multitrait-multimethod analyses of the Big Five in Spanish and English. Journal of Personality and Social Psychology, 75, 729–750.

90. Gosling, S. D., Rentfrow, P. J., & Swann, W. B., Jr. (2003). A Very Brief Measure of the Big Five Personality Domains. Journal of Research in Personality, 37, 504–528.

91. John, O. P & Srivastava, S. (1999). The Big Five trait taxonomy: History, measurement, and theoretical perspectives. In L. A. Pervin, & O. P. John (Eds.), Handbook of personality: Theory and research(pp.102–138). New York: Guilford Press.

92. Kepecs, A., Uchida, N., Zariwala, H. et al. (2008) Neural correlates, computation and behavioural impact of decision confidence. Nature 455, 227–231. https://doi.org/10.1038/nature07200

93. Kotov RI, Bellman SB, Watson DB. Multidimensional Iowa Suggestibility Scale (MISS) Brief Manual[Internet]. 2004 [cited 1 Apr 2015] pp. 7–10. Available: http://medicine.stonybrookmedicine.edu/system/files/MISSBriefManual.pdf

94. Storme M., Tavani J., Myszkowski N., Psychometric Properties of the French Ten-Item Personality Inventory (TIPI) MartinJournal of Individual Differences 2016; Vol. 37(2):81–87.

